# Flanking DNA sequences determine DNA methylation maintenance in proliferation, cancer and aging

**DOI:** 10.64898/2026.04.09.717557

**Authors:** Isaac F. López-Moyado, Lot Hernández-Espinosa, J. Carlos Angel, Aurélie Modat, Ermira Lleshi, Robert Crawford, Geoffrey J. Faulkner, Anjana Rao

## Abstract

DNA methylation is a stable epigenetic modification essential for promoter silencing, retrotransposon silencing, genomic imprinting, and X-chromosome inactivation. Symmetrical DNA methylation at CpG dinucleotides is maintained after every round of cell division by the DNMT1-UHRF1 maintenance methyltransferase complex. Here we define a conserved rank order of DNA hexanucleotide sequences surrounding CpG sites that determines baseline DNA methylation levels in cells and the probability that DNA methylation is retained across cell divisions. This rank order is conserved in vertebrates and does not depend on TET enzymatic activity. CpG sites in hexanucleotide sequences less favored by DNMT1 are more susceptible to replication-dependent loss of DNA methylation over time; consequently, the methylation status of these motifs serves as a marker of cumulative cell divisions, biological age and cancer progression. Thus, the intrinsic vulnerability stemming from the sequence preference of the DNMT1-UHRF1 complex compromises the long-term stability of DNA methylation, especially at heterochromatic sites in proliferating cells, and contributes to the epigenetic dysregulation observed in cancer and aging.

## Introduction

DNA methylation is a fundamental epigenetic modification that plays a critical role in regulating gene expression, maintaining genome stability, and preserving cellular identity (Smith et al., 2025). In mammals, cytosine methylation occurs predominantly at CpG dinucleotides and is dynamically regulated through the opposing actions of DNA methyltransferases (DNMTs), which establish and maintain methylation, and the ten-eleven translocation (TET) family of dioxygenases, which mediate active demethylation. Together, these enzymes ensure the proper balance of DNA methylation at different genomic regions, maintain methylation fidelity, and allow cells to maintain lineage-specific methylation landscapes while still enabling developmental and environmental responsiveness.

The maintenance DNA methyltransferase DNMT1 and its partner UHRF1 preferentially target hemimethylated CpG sites immediately after DNA replication (Bostick et al., 2007; Sharif et al., 2007). These enzymes ensure that methylation patterns are faithfully propagated to daughter strands, preserving epigenetic memory across generations of cells. However, DNA methylation is not perfectly stable over time: cells gradually lose methylation with cell proliferation and aging, even in the absence of obvious perturbations to DNMT or TET enzymes. The molecular determinants of this gradual erosion remain incompletely understood, and whether it reflects an active demethylation process or a passive failure of maintenance methylation is a subject of ongoing debate (Zhou & Reizel, 2025).

Widespread loss of DNA methylation in the genome is a known hallmark of cancer, aging, and senescence (Baylin & Jones, 2016; Cruickshanks et al., 2013; Zhou et al., 2018). In all these conditions, the decrease in DNA methylation occurs predominantly at heterochromatic regions of the genome: for example, the partially methylated domains (PMDs) documented in cancer genomes overlap with late-replicating, nuclear lamina-associated domains enriched for histone marks H3K9me2/H3K9me3 (Berman et al., 2012; Hon et al., 2012; Zhou et al., 2018), and primary human fibroblasts in culture lose DNA methylation in late-replicating regions in a progressive manner during serial passaging (Endicott et al., 2022). The occurrence of these large, megabase-scale, hypomethylated regions of the genome contrasts with the focal increases in DNA methylation which are simultaneously observed in cancer and aging (Moqri et al., 2024); these largely occur at euchromatin sites including cis-regulatory regions such as transcriptional enhancers and promoters (Berman et al., 2012; Rakyan et al., 2010).

TET enzymes oxidize 5-methylcytosine (5mC) to 5-hydroxymethylcytosine (5hmC) (Tahiliani et al., 2009) and further oxidized products (5-formylcytosine, 5-carboxylcytosine) (Ito et al., 2011), contributing to DNA demethylation and the formation of new DNA modifications (López-Moyado et al., 2024). *TET* loss-of-function is commonly observed in cancer; for instance, *TET2* is frequently mutated across hematologic malignancies (Lio et al., 2019). While TET deficiency is typically associated with focal gains of DNA methylation at promoters and enhancers, we previously reported a paradoxical and reproducible loss of DNA methylation across multiple TET-deficient cell types, including somatic cells and embryonic stem cells (López-Moyado et al., 2019). Notably, this decrease was largely observed at heterochromatic regions of the genome, which normally lack both 5hmC (the first product of TET-mediated demethylation) and TETs (López-Moyado et al., 2019).

In this study, we investigated DNA methylation dynamics following *TET* gene deletion to dissect the contributions of TET-dependent and -independent mechanisms to DNA methylation maintenance. By analyzing genome-wide methylation patterns across diverse cell types with distinct proliferative capacities, we show that the DNMT1-UHRF1 complex determines the time-course of loss of DNA methylation at sequences preferred or disfavored by DNMT1 in wildtype and TET-deficient cells. This mechanism is tightly linked to cell division, occurring more rapidly in dividing cells such as lymphocytes compared to non-dividing, post-mitotic neurons. The same rank order of sequence preferences applies in aging and cancer across multiple cell types, suggesting approaches to predicting the time-course of biological aging and cancer progression, as well as investigating the mechanistic basis of both pathological states.

## Results

### TET-independent loss of DNA methylation preferentially affects cytosines in CpG-poor regions

We previously documented a striking increase in proliferation of iNKT cells from *Tet2/3* DKO *CD4Cre* mice (López-Moyado et al., 2019). iNKT cells are generated in the thymus and respond to stimulation with lipid antigens and the noncanonical MHC protein CD1d (Engel & Kronenberg, 2012); however their expansion is severely limited and they normally constitute only a few percent of all T cells in mice. This brake on iNKT cell proliferation observed in normal cells is lost in iNKT cells from *CD4Cre Tet2/3* double-knockout (DKO) cells (López-Moyado et al., 2019), which expand to 10 times their normal levels in spleen and thymus of young (4-5 week-old) mice (Tsagaratou et al., 2017). Even stronger (effectively uncontrollable) proliferation was observed when small numbers of cells from young mice *Tet2/3* DKO *CD4Cre* mice were transferred into wildtype recipient mice (López-Moyado et al., 2019).

Whole-genome bisulfite sequencing revealed an unexpected decrease in heterochromatic DNA methylation in iNKT cells from *Tet2/3* DKO *CD4Cre* mice (López-Moyado et al., 2019). We used Hi-C PC1 values to assign each CpG sufficiently covered by sequencing (*see Methods*) to euchromatic (A compartment; positive PC1 values) or heterochromatic (B compartment; negative PC1 values) regions of the iNKT cell genome (López-Moyado et al., 2019). The loss of DNA methylation in heterochromatin of *Tet2/3* DKO iNKT cells was influenced by cell proliferation, as cells that had been transferred and expanded in a wildtype recipient displayed larger loses of DNA methylation than *Tet2/3* DKO cells from the same mice before adoptive transfer and secondary expansion (López-Moyado et al., 2019). iNKT cells express TET1 mRNA at very low levels (Tsagaratou et al., 2017), suggesting that proliferation and heterochromatic DNA demethylation were unlikely to be due to residual TET1 activity. We therefore hypothesized that the observed loss of DNA methylation upon *TET* deletion might occur through a TET-independent mechanism.

To test this hypothesis directly, we analyzed DNA methylation data from *Tet* triple-knockout (TKO) myeloid cells (Yuita et al., 2023), in which DNA methylation changes should be independent of TET enzymatic activity. Genome-wide DNA methylation analysis revealed that *Tet TKO* myeloid cells broadly recapitulated the previously observed patterns of DNA methylation gain in euchromatin and loss in heterochromatin (**Fig. 1a**). Notably, this loss of DNA methylation was particularly pronounced at CpGs within the solo WCGW sequence context (where W (weak) = A or T; **Fig. 1a**). Comparative analysis across CpG sequence contexts revealed a gradient of DNA methylation loss, with WCGW (^A^/_T_CG^A^/_T_) sites exhibiting the strongest effect, followed by progressively weaker losses in other contexts, such as CpGs flanked by C or G (denoted by S (strong)) (**Suppl. Fig. S1a**) indicating that local nucleotide sequence strongly influences susceptibility to demethylation as noted before (Zhou et al., 2018).

**Figure 1:**
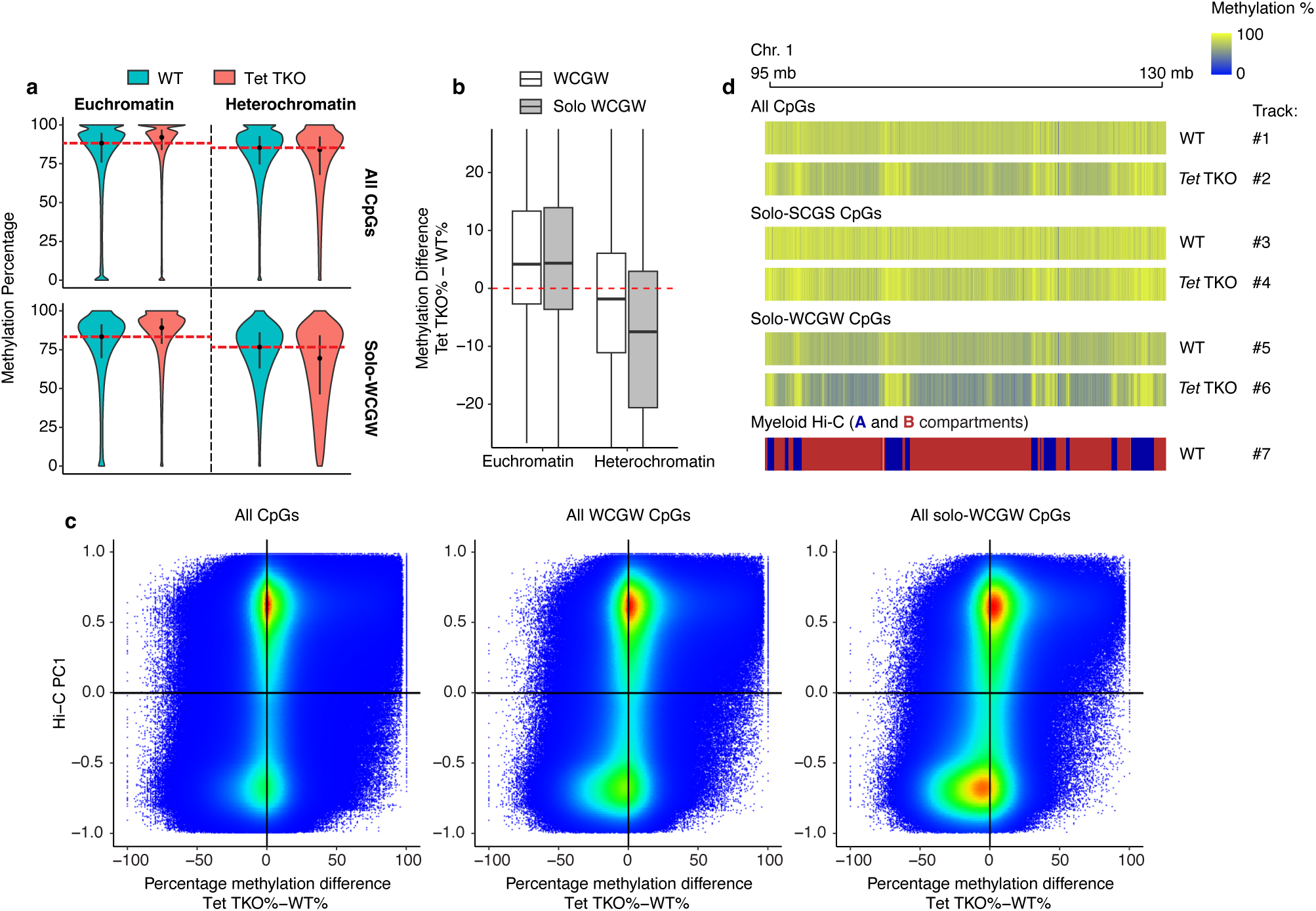
Heterochromatic, sequence-specific, loss of DNA methylation even in fully TET-deficient cells. **(a, b)** DNA methylation percentage (**a**) and methylation difference (**b**) in *Tet1/2/3 TKO* myeloid cells (isolated ∼70 days post-transplantation of hematopoietic progenitors subjected to lentiviral Cre-mediated *Tet* deletion), across all CpGs or CpGs in the solo-WCGW sequence context, in euchromatin and heterochromatin compartments (positive and negative PC1 values in Hi-C). **(c)** Scatterplot colored by density showing the correlation between euchromatin and heterochromatin compartments (y-axis, Hi-C PC1 values) and the difference in DNA methylation percentage between *Tet1/2/3 TKO* myeloid cells and control cells (x-axis) at individual CpGs in different contexts (all CpGs, WCGW, solo-WCGW). Positive Hi-C PC1 values represent euchromatin, negative values represent heterochromatin. Each dot represents an individual CpG. **(d)** Genome tracks showing the DNA methylation percentage in WT versus *Tet1/2/3 TKO* myeloid cells of CpGs in different contexts: All CpGs (tracks 1-2), solo-SCGS CpGs (tracks 3-4) and solo-WCGW CpGs (tracks 5-6), in Hi-C-defined compartments for WT myeloid cells (track 7).

In addition to sequence context, local CpG density emerged as a critical determinant of DNA methylation status. Loss of DNA methylation was significantly more prevalent at CpGs in CpG-poor regions (*solo* CpGs, with no other CpG in a +/- 35 bp window) than at non-solo CpGs, even when analyzing CpGs within the same WCGW sequence context (**Fig. 1b,c**). This observation is consistent with prior reports demonstrating that CpG density influences maintenance methylation efficiency, and supports a model in which isolated CpGs are particularly vulnerable to replication-associated methylation loss (Zhou et al., 2018). Because this phenomenon also occurs in *Tet TKO* cells devoid of TET activity, we conclude that passive, replication-associated DNA demethylation does not require TET activity.

Notably, CpGs across different sequence contexts exhibited highly overlapping distributions within heterochromatin (**Fig. 1c,d**), with solo-WCGW and solo-SCGS CpGs being extensively intermingled. This indicates that, despite their spatial overlap, the solo-WCGW and solo-SCGS sequence contexts are subject to distinct, sequence-dependent mechanisms that give rise to their differential DNA methylation patterns (**Fig. 1d**). Finally, our results are agnostic to the technology used to detect DNA methylation, as similar results were obtained when using WGBS, long-read Oxford Nanopore sequencing, or biomodal 6-base sequencing which detects 5mC and 5hmC in addition to the canonical bases A, C, G, T (**Suppl. Fig. S1b-h**). Thus, the patterns we observe are not attributable to platform-specific biases.

### The hexanucleotide sequence context surrounding a CpG determines its susceptibility to loss of DNA methylation

We previously reported that the heterochromatic DNA methylation loss that occurs upon *Tet2/3* deletion in T cells is influenced by cell proliferation, as iNKT cells from 4-5 week-old *Tet2/3* DKO *CD4Cre* mice that had been transferred and expanded in a secondary recipient displayed larger losses of DNA methylation than *Tet2/3* DKO cells from the same mice before adoptive transfer and secondary expansion (López-Moyado et al., 2019). We analyzed how sequence preference and CpG density influenced DNA methylation dynamics at these two distinct stages of cellular expansion in TET-deficient cells (**Fig. 2a**). We observed that despite their assignment to the same class of sequences, solo-WCGW CpGs in heterochromatin exhibited substantial heterogeneity in the extent of DNA methylation loss (**Fig. 2a**, *see red boxes*). While some CpGs lost up to ∼50% of their DNA methylation upon *Tet2/3* deletion, others retained high methylation levels; this divergence occurred even at similarly negative PC1 values obtained by Hi-C, indicating an equivalent probability of being located in heterochromatin. We used Hi-C PC1 values for iNKT cells (López-Moyado et al., 2019) to assign each CpG sufficiently covered by sequencing to euchromatic (A compartment; positive PC1 values) or heterochromatic (B compartment; negative PC1 values) regions of the genome.

**Figure 2:**
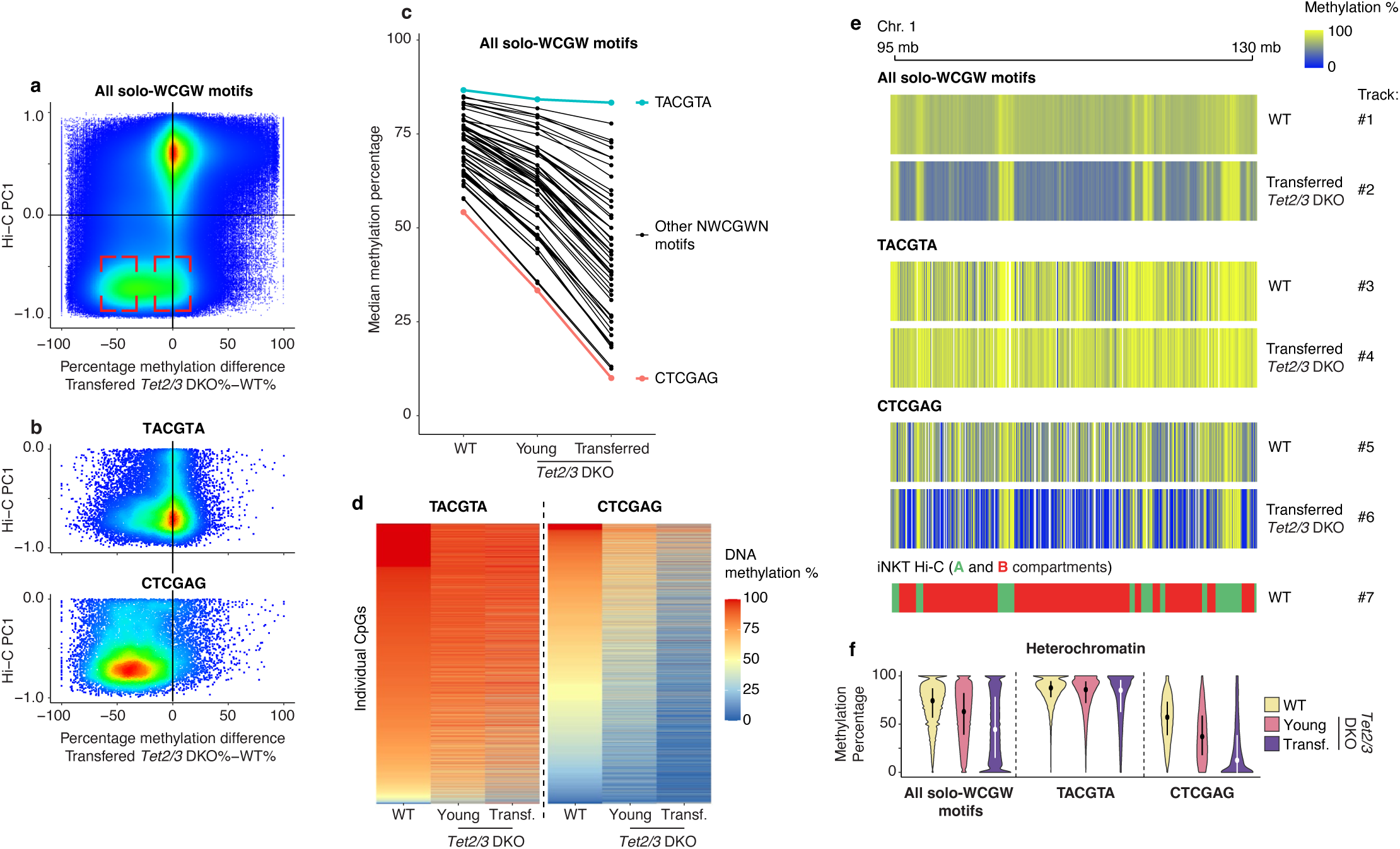
The hexanucleotide sequence context surrounding a CpG determines its DNA methylation level and susceptibility to lose DNA methylation. **(a)** Scatterplot colored by density showing the correlation between euchromatin/heterochromatin compartments (y-axis, Hi-C PC1 values) and the difference in DNA methylation percentage between *Tet2/3 DKO* transferred iNKT cells and control iNKT cells (x-axis) at individual CpGs in the solo-WCGW context. **(b)** Scatterplot colored by density showing the correlation between heterochromatin and the difference in DNA methylation percentage between *Tet2/3 DKO* transferred iNKT cells and controls (x-axis) in solo-CpGs in the TACGTA and CTCGAG sequence contexts. **(c)** Median DNA methylation percentage in WT, *Tet2/3 DKO* young, and *Tet2/3 DKO* transferred iNKT cells, across all 64 possible hexanucleotide CpG sequences found in the solo-WCGW context in heterochromatin. TACGTA and CTCGAG sequences are highlighted in blue and orange, respectively. **(d)** Heatmap of DNA methylation percentage of heterochromatic solo-CpGs in the TACGTA or CTCGAG sequence contexts, in WT, *Tet2/3 DKO* young, and *Tet2/3 DKO* transferred iNKT cells. CpGs are shown in descending order by DNA methylation levels in WT cells. **(e)** Genome tracks showing the DNA methylation percentage in WT and *Tet2/3 DKO* transferred iNKT cells of CpGs in different contexts: All WCGW (tracks 1-2), only TACGTA (tracks 3-4) and only CTCGAG (tracks 5-6), contrasted with the Hi-C-defined compartments for WT iNKT cells (track 7). **(f)** DNA methylation percentage of heterochromatic (Hi-C PC1 value < 0) solo-WCGW CpGs in WT, *Tet2/3 DKO* young, and *Tet2/3 DKO* transferred iNKT cells. Violin plots represent: *left*, all solo-WCGW CpGs; *middle*, solo-WCGW CpGs in the TACGTA context; *right*, solo-WCGW CpGs in the CTCGAG context.

To explore the basis of this heterogeneity, we expanded our analysis by subdividing solo-WCGW CpGs according to the identity of a single additional flanking nucleotide on each side of the core solo-^A^/_T_CG^A^/_T_ motif. This 6 base-pair sequence—hereafter referred to as the *hexanucleotide sequence context*—is defined as the CpG dinucleotide plus the immediately flanking 5’ and 3’ dinucleotides. Across all 64 possible hexanucleotide contexts within heterochromatic solo-WCGW CpGs (comprising 1,253,515 CpGs), we observed marked differences in both DNA methylation levels and susceptibility to demethylation upon *Tet2/3* deletion and cell proliferation (**Fig. 2b,c**; **Suppl. Fig. S2a**). The differences were evident when comparing WT, *Tet2/3* DKO young, and *Tet2/3* DKO transferred iNKT cells, indicating that sequence context modulates DNA methylation dynamics in a proliferation-dependent manner (**Fig. 2b,c**; **Suppl. Fig. S2a**). Notably, CpGs within the TACGTA and CTCGAG hexanucleotide contexts—both of which conform to the WCGW definition—displayed dramatically different behaviors. TACGTA CpGs (comprising 26,921 CpGs) retained high DNA methylation levels upon *Tet2/3* gene deletion regardless of proliferation status, with median methylation decreasing only modestly from 86.6% in WT cells to 83.3% in transferred *Tet2/3* DKO iNKT cells (**Fig. 2b-f**; **Suppl. Fig. S2b-e**). In contrast, CTCGAG CpGs (comprising 17,527 CpGs) exhibited profound DNA methylation loss, dropping from a median of 54.1% in WT cells to approximately 10% in transferred *Tet2/3* DKO cells (**Fig. 2b-f**; **Suppl. Fig. S2b-e**). Importantly, these patterns were consistent across individual CpGs within each hexanucleotide context (**Fig. 2b,d**; **Suppl. Fig. S2b-e**), rather than being driven by a small subset of outliers.

Visualization of DNA methylation patterns along genomic loci further confirmed that CpGs in these distinct hexanucleotide contexts behave differently despite residing within the same heterochromatic regions (**Fig. 2e**). Expanding the sequence window to seven or eight nucleotides (3 or 4 nucleotides flanking the central CG) did not further stratify CpGs by their methylation behavior (**Suppl. Fig. S2b**), indicating that the two nucleotides immediately flanking the CpG are enough to capture the relevant hexanucleotide sequence-dependent effects. Thus, beyond the WCGW classification, the 5’ and 3’ dinucleotides flanking the central CG are critical determinants of both steady-state DNA methylation levels and vulnerability to DNA methylation loss in proliferating *TET*-deficient cells. Our analysis reveals an additional layer of sequence-encoded regulation of DNA methylation maintenance in heterochromatin, which helps explain the pronounced heterogeneity observed among heterochromatic CpGs even within the same solo-WCGW category.

### A similar hierarchy of flanking dinucleotides determines CpG methylation status in both heterochromatin and euchromatin

To dissect in more detail how the hexanucleotide sequence context surrounding a CpG influences DNA methylation across the genome beyond the WCGW sequence context, we analyzed all 15 possible dinucleotide combinations immediately 5’ or 3’ of the CpG dyad in heterochromatic solo CpGs. We excluded flanking sequences containing CGs (i.e., NNCGCG, CGCGNN, CGCGCG) to avoid the confounding effects of tandem CGs in regions of high CpG density, such as CpG islands. When CpGs were grouped by the dinucleotide immediately 5’ of the CpG dyad, we observed a clear hierarchy in DNA methylation levels, with each dinucleotide flanking the central CG associated with a distinct median methylation percentage for the cytosine in the CpG (**Fig. 3a**). An analogous hierarchy was observed for dinucleotides 3’ of the CpG dyad, whose ranking corresponded to the reverse complement of the upstream ranking (**Fig. 3b**). Differences in DNA methylation of the central CpG were better explained by the identity of the 5’ or 3’ dinucleotides flanking the CpG, as opposed to considering the individual flanking nucleotides separately (**Fig. S3a**). Thus, the dinucleotide context, in which the two bases in the dinucleotides flanking the CpG are considered together as a single unit, effectively predicts baseline CpG methylation as well as the extent of methylation decrease after proliferation.

**Figure 3:**
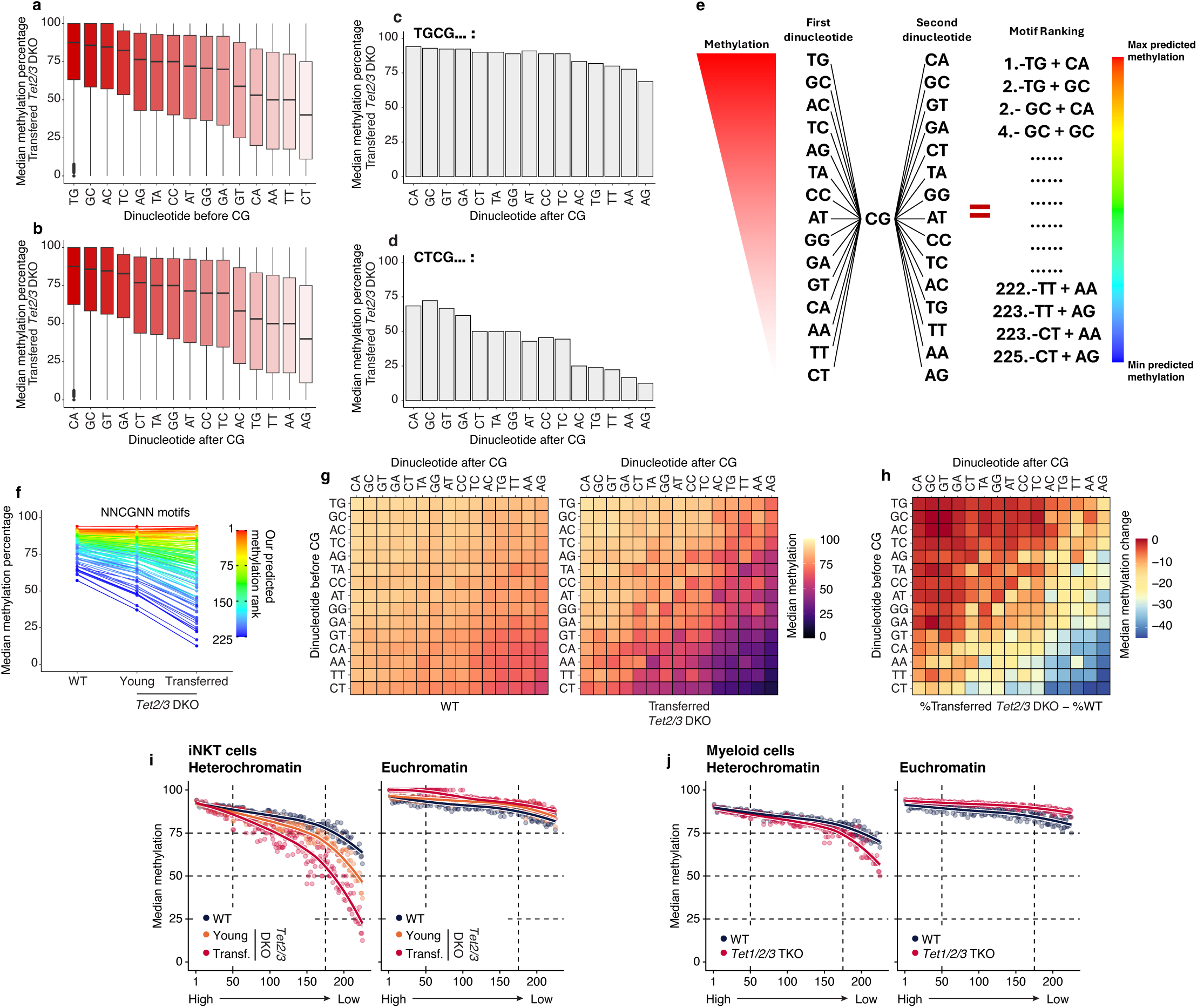
Flanking dinucleotide hierarchy determines the CpG methylation status and dynamics. **(a, b)** DNA methylation percentage of heterochromatic solo-CpGs in *Tet2/3 DKO* transferred iNKT cells, grouped by the dinucleotide before **(a)** or after **(b)** the CG dyad. Dinucleotides are ordered from higher to lower median DNA methylation percentage. **(c, d)** DNA methylation percentage of heterochromatic solo-CpGs in the TGCGNN **(c)** and CTCGNN **(d)** sequence contexts in *Tet2/3 DKO* transferred iNKT cells, grouped by the dinucleotide downstream of the CpG dyad. **(e)** Diagram of the ranking in CpG methylation as determined by a combination of the effects of the upstream and downstream dinucleotides. **(f)** Median DNA methylation percentage in WT, *Tet2/3 DKO* young, and *Tet2/3 DKO* transferred iNKT cells, across all 225 possible hexanucleotide CpG sequences found in the *solo* context in heterochromatin, colored by the predicted methylation rank based on the dinucleotide hierarchy. **(g, h)** DNA methylation percentage at heterochromatic solo-CpGs in WT and *Tet2/3 DKO* transferred iNKT cells **(g),** and difference in CpG methylation between those conditions (*Tet2/3 DKO* transferred% – WT%) **(h)**, displaying all possible combinations of dinucleotides surrounding a CpG (excluding further CG dinucleotides). Dinucleotides are ordered by the predicted methylation rank (highest to lowest rank: top to bottom, left to right). **(i)** Correlation between hexanucleotide CpG motif ranking and median DNA methylation percentage in WT, *Tet2/3 DKO* young, and *Tet2/3 DKO* transferred iNKT cells, for heterochromatic (*left*) and euchromatic (*right*) solo-CpGs. **(j)** Correlation between hexanucleotide CpG motif ranking and median DNA methylation percentage in WT and *Tet1/2/3 TKO* ex vivo myeloid cells, for heterochromatic (*left*) and euchromatic (*right*) solo-CpGs.

We hypothesized that the dinucleotides flanking the CpG dyad exert independent and additive effects on CpG methylation, collectively determining the methylation levels observed across distinct hexanucleotide contexts. To test this, we examined CpGs whose 5’ or 3’ dinucleotides were associated with either high or low median methylation levels. For example, CpGs in the TGCGNN context, which exhibit among the highest methylation levels, and those in the CTCGNN context, which exhibit among the lowest methylation levels, were further stratified by their 3’ dinucleotide (**Fig. 3c, d**). In both cases, the 3’ dinucleotides modulated CpG methylation in a manner that mirrored the 3’ dinucleotide hierarchy identified in **Fig. 3b**, supporting a model in which 5’ and 3’ dinucleotides independently and additively contribute to CpG methylation levels.

Building on this framework, we generated a predictive ranking of all possible 225 hexanucleotide CpG sequences (excluding 31 sequences containing an additional CG dinucleotide flanking the central CpG, to avoid the confounding effects of CpG density, as explained above) by combining the ranks of the upstream (5’) and downstream (3’) dinucleotides (**Fig. 3e**). Each dinucleotide was assigned a rank from 1 to 225 based on its association with CpG methylation levels, and the aggregate of the two flanking dinucleotide ranks defined the predicted methylation rank of the full hexanucleotide sequence (**Fig. 3e**, rainbow gradient at right). These 225 motifs cover 19,612,552 out of the 20,383,623 CpGs in the mm10 genome (96.2% of all the CpGs for mm10), and the dinucleotide-based ranking accurately predicted both steady-state DNA methylation levels and susceptibility to methylation loss across all 225 hexanucleotide contexts in WT, *Tet2/3 DKO* young, and *Tet2/3 DKO* transferred iNKT cells (**Fig. 3f**): sequences with the highest ranking (#1-50) had the highest baseline DNA methylation in WT iNKT cells, and maintained high levels of DNA methylation following proliferation (**Fig. 3e**, *right*, red in the rainbow gradient; **Fig. 3f**). Conversely, sequences with the lowest ranking (#175-225) had the lowest baseline levels of DNA methylation in wildtype iNKT cells, and lost the greatest amount of DNA methylation after proliferation (**Fig. 3e**, *right*, blue in the rainbow gradient; **Fig. 3f**). Interestingly, both the top (#1, TGCGCA) and bottom (#225, CTCGAG) sequences are palindromic, and belong to the SGCS and WGCW sequences contexts respectively. Matrices displaying DNA methylation levels and methylation changes across all dinucleotide combinations revealed a linear and graded effect of each flanking dinucleotide, incorporating the effects of both 5’ and 3’ dinucleotides in the ranking (**Fig. 3g, h**). The strong correlation between predicted methylation rank and observed DNA methylation levels further validated the robustness of this model (**Suppl. Fig. S3a**).

All the analyses described above were initially restricted to heterochromatic CpGs (**Suppl. Fig. S3b,c**), where DNA methylation loss is the most pronounced in *TET*-deficient cells, similar to what occurs in cancer and replication-associated methylation loss. However, applying the same ranking system to euchromatic CpGs revealed a similarly strong correlation between predicted rank and DNA methylation levels, albeit with reduced dynamic range (**Fig. 3i,j**). This indicates that the sequence-encoded determinants of CpG methylation operate broadly across the genome, independent of chromatin compartment. Notably, we observed comparable dinucleotide-driven methylation patterns in *Tet1/2/3* triple-knockout (TKO) myeloid cells (**Fig. 3i**), demonstrating that the sequence-dependent hierarchy that we identify is not specific to iNKT cells, and operates even in the complete absence of TET enzymatic activity. Thus, CpG methylation status and dynamics can be predicted by a hierarchy of flanking 5’ and 3’ dinucleotides that each display a marked periodicity with respect to WS and purine-pyrimidine identities (**Suppl. Fig. S3d**), revealing an intrinsic, sequence-encoded layer of DNA methylation regulation acting genome-wide.

### The hierarchy of flanking dinucleotides predicts the replication-dependent loss of DNA methylation in numerous cellular systems

So far, we have shown that the hexanucleotide hierarchy we have identified predicts the methylation dynamics of nearly all CpGs genome-wide. This was true for profoundly (*Tet2/3 DKO*) as well as completely (*Tet1/2/3 TKO*) TET-deficient cells in at least two primary cell types (T cells and myeloid cells) (**Fig. 3, Supp. Fig. 3**). The dependence on cell proliferation but not TET proteins predicts that the underlying mechanism involves DNA methyltransferase (DNMT) enzymes, and specifically that the likelihood of CpG remethylation by the maintenance DNMT1/UHRF1 complex depends in a predictable fashion on flanking dinucleotides. The sequence preference is most evident in regions where methylation maintenance is inherently less efficient, such as heterochromatic domains, late-replicating regions, and partially methylated domains (PMDs) (Salhab et al., 2018; Zhou et al., 2018) (**Fig. 3**, **Supp. Fig. 3**). We therefore used published whole-genome bisulphite sequencing and other comprehensive genome-wide DNA methylation analyses to investigate dynamic DNA methylation differences in different cell types sampled under different conditions.

Using DNA methylation arrays covering almost a million CpGs, a previous study (Endicott et al., 2022) demonstrated that heterochromatic sequences in human primary fibroblasts from different sources lose DNA methylation upon serial passaging due to incomplete DNA methylation maintenance during successive replications. Our hexanucleotide ranking accurately predicted the DNA methylation changes observed in that study (**Suppl. Fig. S4a**). In each case, low-ranking sequences (#175-225) failed to regain full methylation and progressively lost 5mC marks on both strands (**Suppl. Fig. S4a**).

To extend these observations, we analyzed publicly available methylome datasets for biological contexts associated with replication-coupled DNA methylation loss—aging, cancer and cellular senescence. In each case—T cells from a newborn versus a 103-year-old centenarian (Heyn et al., 2012) (**Fig. 4a**; **Supp. Fig. S4b**); colorectal tumors versus adjacent normal colon tissue (Berman et al., 2012) (**Fig. 4b**), or IMR90 fibroblasts before and after replication-induced senescence (Cruickshanks et al., 2013) (**Fig. 4c**), the hexanucleotide CpG ranking correlated with DNA methylation dynamics. Higher-ranking motifs (#1-50) retained CpG methylation, whereas lower-ranking motifs (#175-225) displayed progressive loss of CpG methylation, with a more pronounced slope in the lowest-ranking motifs. This pattern reinforces the concept that the hexanucleotide flanking sequence determines replication-dependent methylation maintenance in diverse cellular contexts.

**Figure 4:**
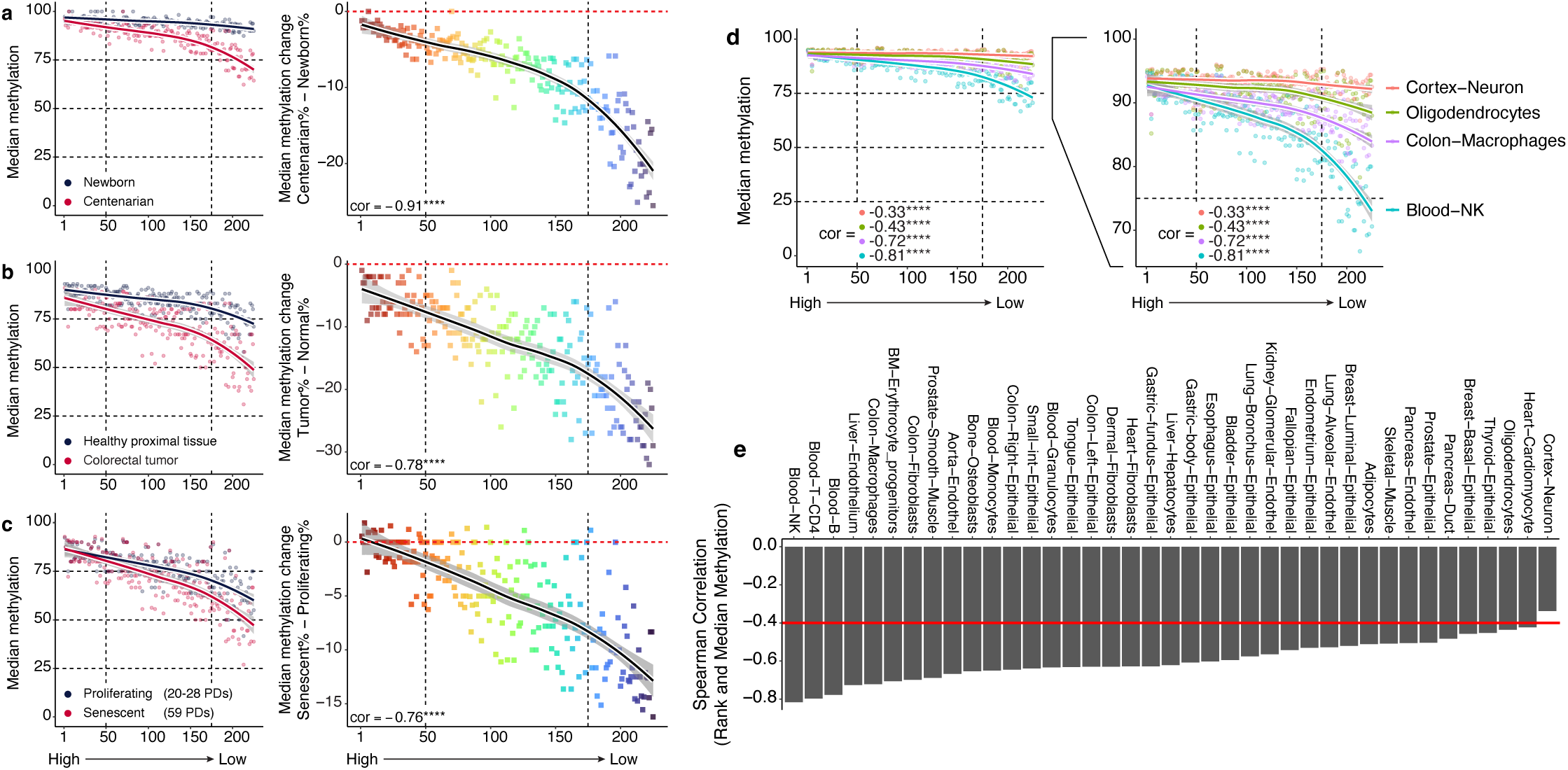
Differential DNA methylation loss observed across hexanucleotide CpG motifs is cell replication-dependent. (**a**-**c**) Genome-wide correlation between hexanucleotide CpG motif ranking and median DNA methylation percentage (*left*), or median DNA methylation difference (*right*), comparing **a**, CD4^+^ T cells from a newborn with those of a centenarian; **b**, colorectal cancer cells with healthy proximal tissue; and **c**, proliferating cells and senescent cells during replication-induced senescence in IMR90 fibroblasts. Fitted regression lines along their confidence intervals are displayed for each scatter plot. Spearman correlation scores for each DNA methylation difference scatter plot are shown in the bottom left corner of each right panel. (**d**) Genome-wide correlation between hexanucleotide CpG motif ranking and median DNA methylation percentage in two primarily post-mitotic cell types (cortex neurons and oligodendrocytes), and two rapidly proliferating cell types (Natural Killer cells and colon macrophages). Fitted regression lines along their confidence intervals plus Spearman correlation scores are displayed for each scatter plot. Notice the association between proliferation rate and the magnitude of correlation score. (**e**) Spearman correlation scores obtained by comparison of hexanucleotide CpG motif ranking and median DNA methylation percentages from 39 diverse cell types across the human body. Cell types ordered by the magnitude of correlation (where a correlation score closer to -1 indicates a strong relationship between DNA methylation values and the proposed hexanucleotide CpG motif rankings). **** denotes *p* < 0.0001 (Spearman correlation).

DNA methylation levels consistently correlated with hexanucleotide rankings across a broad range of 205 human cell types (Loyfer et al., 2023, human DNA methylation atlas), although the magnitude of correlation varied widely (Spearman’s ρ = -0.82 to -0.34) (**Fig. 4d,e**; **Suppl. Fig. S4c**). Rapidly proliferating cells, such as blood lineages, displayed the strongest negative correlations, whereas post-mitotic or terminally differentiated cells (neurons, adipocytes, cardiomyocytes, skeletal muscle) showed weaker correlations (**Fig. 4e**; **Suppl. Fig. S4c**). *Tet1/2/3*-deficient retinal cells, which are largely post-mitotic (Dvoriantchikova et al., 2025), did not show a strong correlation of loss of DNA methylation with the hexanucleotide sequence ranking. Rather, they showed a minimal drop in DNA methylation levels, from ∼93% to ∼83% for the motif that lost the most methylation (**Suppl. Fig. S4d**), compared to TET-deficient myeloid and iNKT cells in which the extent of reduction of heterochromatic DNA methylation in the lowest-ranking motif (#225) was much greater (∼90% to ∼25% in transferred and expanded iNKT cells, and from ∼85% to ∼45% in myeloid cells) (**Fig. 3i,j**; **Suppl. Fig. S3b,c**). Methylation differences across cell types primarily arose from variation in low-ranking motifs, whereas high-ranking motifs remained largely invariant, consistent with their efficient maintenance across replication events (**Suppl. Fig. S4c**).

### Loss of DNA methylation at low-ranking CpG motifs provides an estimate of biological age

Given the strong association between loss of DNA methylation at specific hexanucleotide CpG motifs and cell proliferation (**Suppl. Fig. S4c**), we reasoned that this motif-specific behavior would also be closely linked to aging. In particular, we reasoned that certain hexanucleotide CpG motifs are intrinsically prone to retain DNA methylation during repeated cell divisions, whereas others preferentially lose methylation as cells proliferate, and the latter could be used to provide a quantitative readout of cumulative proliferative history and, by extension, biological age. Since DNA methylation loss is most pronounced in lymphocytes (blood B, CD4^+^ T and NK cells, **Fig. 4e**; iNKT cells, **Fig. 3i**), we focused our analysis on aging in mouse and human T cells.

We first reanalyzed whole-genome DNA methylation profiles that had been used to define the epigenetic features of physiological aging during prolonged, cell-autonomous proliferation of mouse CD8⁺ T cells (Mi et al., 2024). WGBS was performed on naïve CD8⁺ T cells isolated from aged mice (∼2 years old), antigen-specific “young” memory CD8⁺ T cells (∼180 days post-immunization), and memory CD8⁺ T cells that had undergone repeated cycles of adoptive transfer into congenically distinct naïve hosts and repeated antigenic stimulation. These adoptively transferred memory cells were maintained for ∼0.5 lifetimes (∼450 days), 2 lifetimes (∼1350–1620 days), and up to 4 lifetimes (∼2880 days), thereby generating T cells whose proliferative history far exceeded the normal murine lifespan (Mi et al., 2024). Consistent with prior reports that naïve T cells appear epigenetically “young” regardless of host age (Mi et al., 2024), we observed uniformly high DNA methylation levels across the full range of hexanucleotide motif ranking in naïve CD8⁺ T cells (**Fig. 5a**, *yellow line*). In contrast, memory CD8⁺ T cells exhibited a progressive, proliferation-dependent loss of DNA methylation, particularly at low-ranking CpG motifs (**Fig. 5a**, *orange line*). Specifically, high-ranking motifs (#1-50) remained largely stable across memory cells that had proliferated for 0.5 to 4 mouse lifetimes, whereas low-ranking motifs (#175-225) displayed increasing methylation loss that closely tracked chronologic time (**Fig. 5a**, *red to black lines*), with an almost linear relationship between the magnitude of DNA methylation loss at each CpG motif and our hexanucleotide ranking **(Fig. 5b**). Analysis of individual motifs further illustrated this effect: among the top 10 high-ranking motifs (#1-10), DNA methylation levels remained consistently high (>90%) across all proliferative states, whereas among the bottom 10 low-ranking motifs (#216-225), DNA methylation progressively declined from ∼90% in naïve cells, to ∼60% in young memory cells, and to ∼25% in memory cells that had proliferated for 4 lifetimes (**Fig. 5c**).

**Figure 5:**
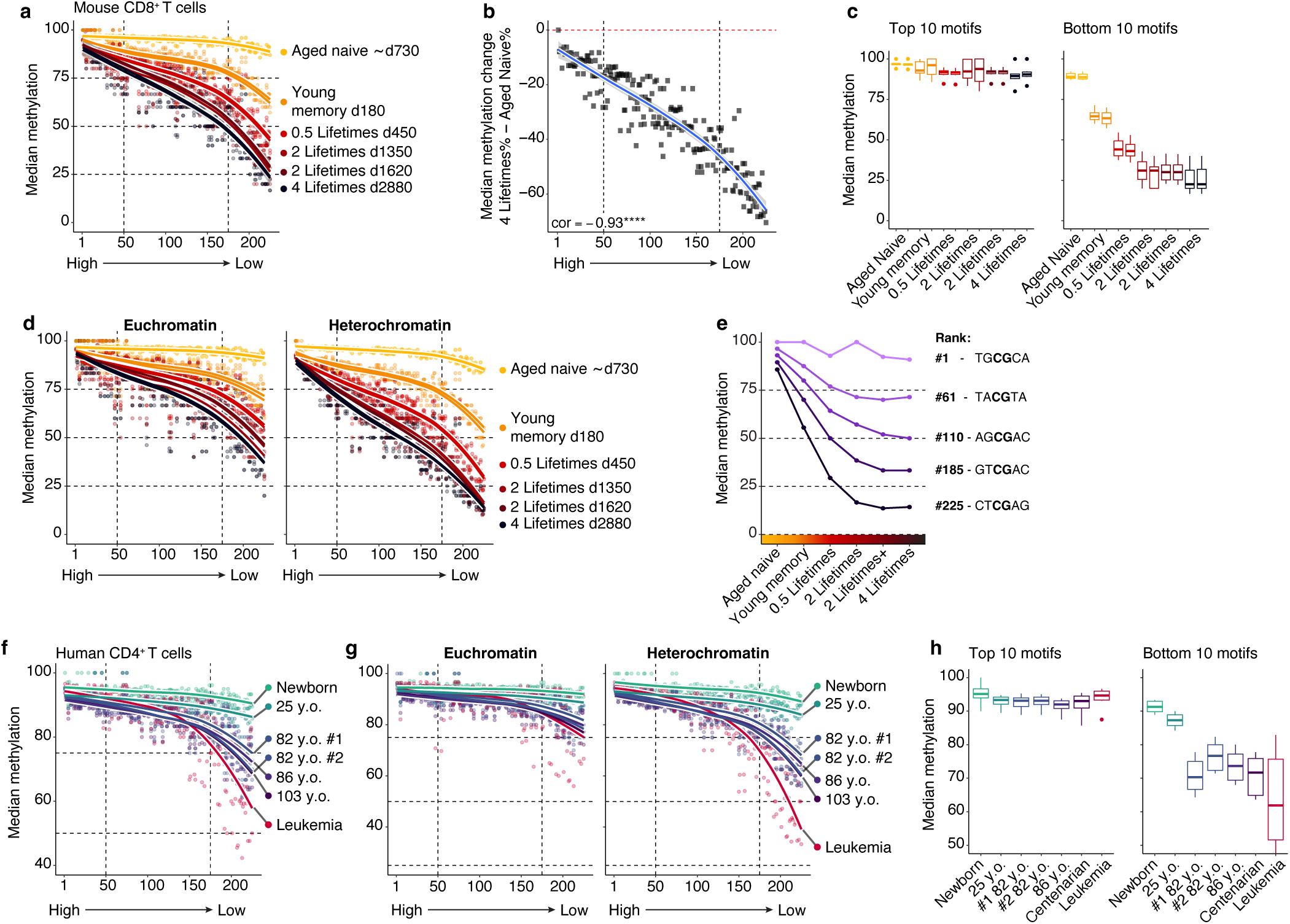
Loss of DNA methylation at low-ranking CpG motifs provides an estimate of cell proliferation magnitude and biological age. (**a**) Genome-wide correlation between hexanucleotide CpG motif ranking and median DNA methylation percentage in different subsets of mouse CD8+ T cells, comparing naïve CD8+ T cells from ∼730-day-old (∼2- year-old) mice to CD8+ T cells that were stimulated and transferred serially into recipient mice, thereby achieving the indicated cumulative lifespans. (**b**) Median DNA methylation difference, comparing aged naïve CD8 T cells with those that have been transferred for 4 lifetimes (from (**a**)). Fitted regression lines along their confidence intervals are displayed for each scatter plot. Spearman correlation scores for each DNA methylation difference scatter plot are shown (*bottom left corner*). **** denotes *p* < 0.0001 (Spearman correlation). (**c**) Median DNA methylation percentage for the top 10 hexanucleotide motifs (ranked 1-10) and bottom 10 motifs (ranked 216-225), across different subsets of mouse CD8+ T cells (from (**a**)). (**d**) Genome-wide correlation between hexanucleotide CpG motif ranking and median DNA methylation percentage in different subsets of mouse CD8+ T cells (from **a**), dividing CpGs by their location in euchromatin vs heterochromatin. The difference in slopes at different ages is particularly prominent in heterochromatin. (**e**) Change in DNA methylation in heterochromatin for five different hexanucleotide motifs (spanning different ranks), in mouse CD8+ T cells proliferating for different mouse lifespans (from (**d**), *right panel*). (**f**) Genome-wide correlation between hexanucleotide CpG motif ranking and median DNA methylation percentage in human CD4+ T cells across donors from different ages (newborn-100 years old). Data from human T cell leukemia cells is shown for comparison. (**g**) Data from (**f**), classifying CpGs by their location in euchromatin vs heterochromatin. (**h**) Median DNA methylation percentage for the top 10 sequences/motifs (ranked 1-10) and bottom 10 sequences (ranked 216-225) in human CD4+ T cells across donors from different ages.

Stratification of CpGs by chromatin context revealed that while hexanucleotide motif-associated methylation loss occurred in both euchromatin and heterochromatin, the effect was particularly pronounced in heterochromatin and in memory CD8⁺ T cells that had proliferated for 4 mouse lifetimes (**Fig. 5d**, *compare black lines*). In heterochromatic CpGs, analysis of representative hexanucleotide CpG motifs spanning five different motif ranks demonstrated progressive, proliferation-dependent methylation loss with increasing cell age and chronologic time (**Fig. 5e**). The highest-ranking motifs (#1, TGCGCA) showed stable methylation across all proliferative states, whereas progressively lower-ranking motifs exhibited increasing methylation loss; methylation in the lowest-ranking motifs (#225, CTCGAG) declined from ∼85% in naïve cells to ∼15% in 4-lifetime cells. Methylation loss at these heterochromatic CpGs appeared to plateau between 2 and 4 lifetimes, suggesting a lower bound for each sequence motif below which methylation is not further lost (**Fig 5e**). This limit depends on the precise sequence of each motif, suggesting a steady state for each motif maintained by the TET-DNMT1 balance, the DNMT1-de novo DNMT3A/DNMT3B balance, or both.

Loss of DNA methylation at low-ranking motifs also tracked with T cell proliferation in other biological contexts. In a mouse LCMV infection model, effector CD8⁺ T cells exhibited greater methylation loss at low-ranking motifs than acute memory or naïve CD8⁺ T cells, and sustained proliferation during chronic infection further accentuated this effect (**Suppl. Fig. S5a,b**). Similarly, CD8⁺ T cells subjected to PD-1 blockade during chronic infection showed slightly enhanced methylation loss at low-ranking motifs, consistent with increased proliferative activity (**Suppl. Fig. S5a,b**).

### Conservation of hexanucleotide CpG motif ranking across vertebrates

To determine whether this cell-autonomous, motif-specific DNA methylation loss is conserved in humans and associated with organismal aging, we reanalyzed DNA methylation data from human CD4⁺ T cells isolated from donors spanning a broad age range, including newborns, young adults (∼25 years), elderly individuals (80–100 years) (Heyn et al., 2012; Jenkinson et al., 2017), and a T cell leukemia sample for reference (Adams et al., 2012). High-ranking motifs (#1-100) showed very little change; low-ranking motifs (#150-225) displayed progressive methylation loss with increasing donor age; and leukemic T cells exhibited even more pronounced methylation loss consistent with rapid proliferation (**Fig. 5f, g**). The extent of methylation loss in the leukemia was considerably more variable than observed in aged cells, consistent with varying proliferation rates in a heterogeneous cell population with undergoing clonal selection (**Fig. 5f**). Further analysis confirmed that low-ranking hexanucleotide motifs preferentially lost DNA methylation with increasing age, whereas high-ranking motifs remained comparatively stable across the human lifespan (**Fig. 5h**). As observed in mice, stratification by chromatin context revealed that age-associated methylation loss at low-ranking motifs was most pronounced in heterochromatin, where motif ranking strongly predicted DNA methylation levels across age groups (**Fig. 5g**).

To explore the cross-species conservation of our hexanucleotide sequence ranking, we reanalyzed whole-genome DNA methylation profiles from fibroblasts of multiple vertebrate species (including human, mouse, rabbit, dog, cow, pig and chicken) (Al Adhami et al., 2022), as well as cells from *Arabidopsis thaliana* (Williams et al., 2022). In all the vertebrate species, differences in DNA methylation levels across CpG sequence contexts followed our predicted hexanucleotide motif ranking (**Suppl. Fig. S5c**), but this correlation was not observed in *Arabidopsis* (**Suppl. Fig. S5c**). DNMT1 and its partner UHRF1 are evolutionarily conserved in vertebrates, consistent with a common underlying mechanism shaping DNA methylation patterns in vertebrates that does not extend to the distinct DNA methylation machinery used by plants (Iyer et al., 2011). We conclude that loss of DNA methylation at low-ranking CpG motifs provides a conserved, cross-species estimate of cumulative cell proliferation and biological age in vertebrates.

### DNMT1 activity correlates with CpG hexanucleotide ranking

Because DNA methylation patterns must be faithfully copied during each round of DNA replication, we hypothesized that the sequence-dependent methylation loss observed at low-ranking hexanucleotide motifs reflects imperfect maintenance by the replication-coupled DNA methyltransferase DNMT1. To test this hypothesis, we reanalyzed published *methylation fidelity* data obtained by hairpin bisulfite sequencing in mouse embryonic stem cells (mESCs) before and after differentiation (and hence slower cycling) induced by LIF withdrawal (Zhao et al., 2014). This assay directly quantifies CpGs that are symmetrically methylated, unmethylated, or hemimethylated following DNA replication. Across both undifferentiated and differentiated states, methylation fidelity (quantified as the percentage of symmetrically methylated CpGs in total CpGs) correlated strongly with our CpG hexanucleotide motif ranking (**Suppl. Fig. S6a**), with high-ranking motifs (#1-50) exhibiting greater fidelity and low-ranking motifs (#175-225) showing increased hemimethylation. Although overall methylation fidelity was higher in differentiated cells from which LIF had been withdrawn, the relative hexanucleotide motif ranking was preserved, indicating that CpG-flanking sequence context intrinsically influences the probability that hemimethylated CpGs are correctly remethylated after replication. These data point to a sequence-dependent component of DNMT1-mediated DNA methylation maintenance, likely involving the DNMT1-UHRF1 complex.

To ask whether DNMT1 alone is sufficient to generate the observed hexanucleotide motif ranking, we reanalyzed WGBS data from human embryonic stem cells (hESCs) lacking DNMT3A, DNMT3B, and all three TET enzymes, leaving DNMT1 as the sole DNA methyltransferase (DNMT1-only cells) (Charlton et al., 2020) (**Fig. 6a**). Comparison of early (passage 6) and late (passage 20) cultures in DNMT1-only cells revealed preferential retention of DNA methylation at high-ranking hexanucleotide motifs, whereas low-ranking motifs exhibited significantly greater methylation loss over time. This pattern was evident both in absolute DNA methylation levels and in passage-dependent methylation changes (**Fig. 6a; Suppl. Fig. S6b**), demonstrating that DNMT1-mediated maintenance is intrinsically biased toward sequence contexts containing high-ranking motifs, and is less effective at maintaining DNA methylation at low-ranking motifs during prolonged proliferation.

**Figure 6:**
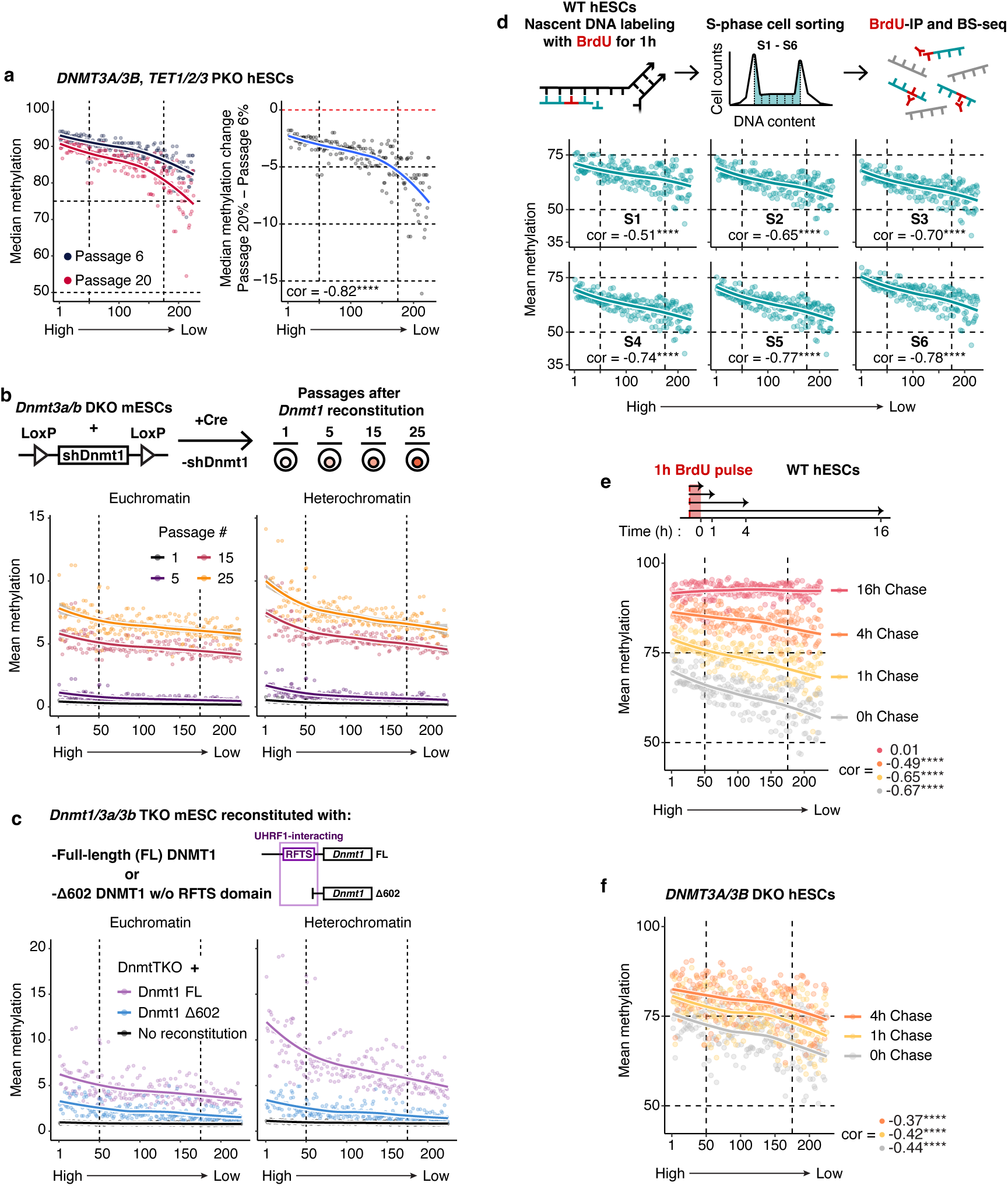
DNMT1 activity is correlated with CpG hexanucleotide ranking. (**a**) Genome-wide correlation between hexanucleotide CpG motif ranking and median DNA methylation percentage in human ESC expressing only DNMT1 (PKO hESC, pentuple knockout: DNMT3A/B DKO and TET1/2/3 TKO). *Left*, data from culture passages 6 and 20; *right*, median DNA methylation difference between passage 6 and passage 20. Spearman correlation score for DNA methylation difference scatter plot is shown (*bottom left corner*). **** denotes *p* < 0.0001 (Spearman correlation). (**b**) WGBS data for *Dnmt3a/3b* mouse ESC, expressing a *floxed* shRNA against *Dnmt1* that can be deleted with Cre to restore DNMT1 protein expression, were analysed at different passages after Cre-mediated deletion and restoration of DNMT1 expression. The genome-wide correlation between hexanucleotide CpG motif ranking and mean DNA methylation percentage is shown. CpGs were classified by their location in euchromatin vs heterochromatin. (**c**) Genome-wide correlation between hexanucleotide CpG motif ranking and mean DNA methylation percentage in *Dnmt TKO* mouse ESC upon reconstitution of DNMT1 protein expression, either with full length (FL) DNMT1 or a truncated version lacking the UHRF1-interacting N-terminal domain (amino acids 1-602, referred as Δ602). CpGs were classified by their location in euchromatin vs heterochromatin. (**d**) Genome-wide correlation between CpG hexanucleotide motif ranking and median DNA methylation levels in human ESCs across sequential S-phase fractions (S1–S6) measured by Repli-BS following BrdU labeling. Cells from later stages of S-phase displayed increased correlation of DNA methylation with motif ranking, indicating progressive DNMT1-mediated remethylation of high-ranking hemi-methylated motifs during the time course of DNA replication. (**e**) DNA methylation levels across CpG hexanucleotide motifs following BrdU pulse–chase labeling in wild-type human ESCs. Early time points exhibit a pronounced methylation difference between high- and low-ranking motifs, which is resolved at later time points, consistent with completion of DNA methylation maintenance and *de novo* DNA methylation. (**f**) DNA methylation levels across CpG hexanucleotide motifs following BrdU pulse–chase labeling in DNMT3A/B double-knockout (DKO) human ESCs. In contrast to wild-type cells, the methylation gap between high- and low-ranking motifs persists over time, indicating preferential DNMT1 activity at high-ranking motifs in the absence of de novo methyltransferases. Fitted regression lines plus Spearman correlation scores for each BrdU pulse-chase Repli-BS median DNA methylation scatter plot are shown (**d**, *bottom*; **e-f**, *bottom right corner*). **** denotes *p* < 0.0001 (Spearman correlation).

To directly assess DNMT1 enzymatic activity, we reanalyzed data from DNMT3A/3B-deficient mESCs in which DNMT1 expression was silenced by an shRNA that could be inducibly excised from the genome, hence restoring DNMT1 expression after Cre recombinase was introduced (Haggerty et al., 2021). This system measures the *de novo* methylation ability of DNMT1 during successive cell divisions after DNMT1 expression was restored. DNA methylation gradually accumulated over multiple passages following Cre expression, reaching 6-10% genome-wide by passage 25. Even this low level of methylation recovery (which reflects the poor de novo methylation ability of DNMT1, (Haggerty et al., 2021)) consistently reached higher levels at high-ranking hexanucleotide motifs than at low-ranking motifs (**Fig. 6b**). The bias was particularly pronounced in heterochromatin, mirroring the chromatin context in which age-associated methylation loss is most evident in T cells (**Fig. 5**) and in transformed *TET*-deficient somatic cells (**Fig. 3**). The preference for heterochromatin strongly suggests the involvement of UHRF1, which recruits DNMT1 to heterochromatin through the ability of its tandem TUDOR domain to recognize the heterochromatic histone modification H3K9me3 (Liu et al., 2013; Rothbart et al., 2012).

To provide further support for UHRF1 involvement, we reanalyzed a study in which DNMT triple-knockout mESCs were reconstituted with either full-length DNMT1 or a truncated DNMT1 lacking the UHRF1-interacting RFTS domain (lacking amino acids 1 to 602; the mutant protein is termed Δ602) (Ito et al., 2024) (**Fig. 6c**). Reconstitution with Δ602 DNMT1 restored only low levels of DNA methylation with modest hexanucleotide motif dependence, but reconstitution with full-length (FL) DNMT1 led to substantially higher methylation gains, particularly in heterochromatin, with a strong preference for high-ranking hexanucleotide motifs. Specifically, full-length DNMT1 restored ∼12% methylation at high-ranking motifs (#1-50) and ∼5.5% at low-ranking motifs (#175-225) in heterochromatin, compared to Δ602 DNMT1 where the corresponding numbers were ∼3.5% (motifs #1-50) and ∼1.5% (motifs #175-225) (**Fig. 6c**). Together, these data indicate that DNMT1 exhibits higher maintenance as well as *de novo* activity at high-ranking hexanucleotide motifs, a property that is enhanced by interaction with UHRF1.

Given the tight coupling between DNMT1 activity and DNA replication, we next examined methylation inheritance dynamics using Repli-BS data from wildtype human ESCs, which combines BrdU labeling with bisulfite sequencing to profile newly replicated DNA (Charlton et al., 2018). CpGs analyzed across six S-phase fractions, defined as S1–S6, recapitulated our proposed hexanucleotide ranking, with later S-phase fractions showing stronger correlations between motif ranking and DNA methylation levels (range -0.51 to -0.78) (**Fig. 6d**). BrdU pulse-chase Repli-BS experiments performed in the same study (Charlton et al., 2018) showed that in wild-type hESCs, initial methylation differences between high- and low-ranking motifs were resolved over time (**Fig. 6e**), confirming that the combined actions of DNMT1 and *de novo* DNMT3A/B enzymes establish DNA methylation genome-wide in hESCs. In contrast, in DNMT3A/B double-knockout hESCs, the methylation gap between high-and low-ranking motifs persisted over time (**Fig. 6f**), further demonstrating preferential DNMT1 activity at high-ranking motifs, and indicating that DNMT1 alone is insufficient to establish DNA methylation across all sequence contexts. Together, these findings indicate that while DNMT1 is capable of establishing and maintaining DNA methylation independently, it does so imperfectly and with clear sequence preferences.

Finally, to distinguish sequence-dependent maintenance inefficiency—the product of imperfect DNMT1 activity—from methylation loss caused by direct DNMT1 inhibition, we analyzed conditions in which DNMT1 activity is inhibited. In mESCs treated with the DNMT1-specific inhibitor GSK-3484862 (Azevedo Portilho et al., 2021), widespread loss of DNA methylation occurred but showed no correlation with hexanucleotide ranking (**Suppl. Fig. S6c**), indicating that the sequence-dependent hexanucleotide ranking requires DNMT1-mediated maintenance. Altogether, our findings establish CpG hexanucleotide motif ranking as a proxy for intrinsic preferential DNMT1 maintenance efficiency, and provide a mechanistic foundation for the progressive, replication-dependent loss of DNA methylation at low-ranking motifs that accumulates with cellular proliferation.

## Discussion

In this study, we identify intrinsic differences among CpG hexanucleotide motifs in their ability to maintain DNA methylation during DNA replication and cell proliferation. By ranking CpG motifs according to their susceptibility to loss of DNA methylation upon extensive proliferation, we show that CpGs embedded in specific hexanucleotide sequence contexts are inherently more prone to DNA hypomethylation whereas others are preferentially maintained, and this difference impacts steady-state DNA methylation levels across CpGs in different hexanucleotide sequence contexts. Analyses across diverse datasets—spanning multiple cell types, organisms, and biological contexts including cancer, aging, and T cell activation—demonstrate that this hexanucleotide CpG hierarchy is evolutionarily conserved, particularly among vertebrates (**Suppl. Fig. S5c**), and universal across cell types. Our findings thus highlight an additional regulatory layer determining DNA methylation levels, beyond established determinants of DNA methylation maintenance, including replication timing (Charlton et al., 2018; Ming et al., 2020), CpG density (Zhou et al., 2018), histone modifications (e.g., H3K9me3) (Ren et al., 2020), and chromatin remodeling (Dennis et al., 2001; Ming et al., 2020). Our results demonstrate that <u>DNA methylation levels are directly influenced by the local hexanucleotide DNA sequence</u>, adding a DNA sequence-encoded dimension to epigenetic regulation.

Our data indicate that differential methylation loss across CpG hexanucleotide motifs is largely replication-dependent and occurs across biological conditions characterized by proliferation, from development and immune activation to aging and cancer. Multiple lines of evidence point to a fundamental property of the DNMT1-UHRF1 “maintenance” DNA methylation complex, which remethylates hemimethylated CpGs in each round of DNA replication (Liu et al., 2013; Ming et al., 2020; Nishiyama et al., 2020; Rothbart et al., 2012), as the central mediator of this effect. For instance, in cellular models in which DNMT1 is the sole active DNA methyltransferase (**Fig. 6**), hexanucleotide motif-specific differences in DNA methylation levels and maintenance activity are observed, indicating that they arise from differential DNMT1 activity rather than from *de novo* methylation or active demethylation. Likewise, this DNMT1-associated motif preference disappears when cells are treated with a specific inhibitor of DNMT1 enzymatic activity (**Supp. Fig. 6c**). Therefore, we propose that CpGs that preferentially lose methylation during extensive proliferation correspond to sequence contexts in which DNMT1-mediated DNA methylation maintenance is intrinsically less efficient.

Mechanistically, our results support a model in which DNMT1-UHRF1-mediated maintenance methylation operates with imperfect, sequence-biased efficiency. DNMT1 restores symmetric CpG methylation following replication through a process requiring cytosine base flipping and conformational rearrangements of the catalytic pocket (Adam et al., 2020), while UHRF1 recognizes hemimethylated DNA via its SRA domain (Hashimoto et al., 2008). Structural and biochemical studies have shown that these steps are sensitive to DNA shape, groove geometry, and local flexibility (Adam et al., 2020). Consistent with this, crystal structures of DNMT1 bound to favored versus disfavored sequence contexts reveal distinct DNA rearrangements and catalytic site conformations that correlate with methylation efficiency (Adam et al., 2020). Since both DNMT1 and UHRF1 engage the major and minor grooves (Hashimoto et al., 2008; Song et al., 2011), it is very plausible that there is a DNA structural basis for how flanking sequence could modulate enzymatic activity. Importantly, the loss of DNA methylation at low-ranking sequences—associated with cell proliferation—is unlikely driven by differences in DNMT1 or UHRF1 expression, whose mRNA and protein levels are regulated in a cell-cycle-dependent way (Du et al., 2010). Instead, we propose that local DNA structure—encoded by the hexanucleotide sequence surrounding a CpG motif—biases DNMT1 activity at individual CpG sites.

Our analysis of diverse *TET*-deficient systems in this study allowed us to separate replication-coupled DNA methylation loss from enzymatic demethylation, since in the absence of TET-mediated enzymatic demethylation, any observed methylation loss must arise from imperfect maintenance rather than active conversion of 5-methylcytosine (Tahiliani et al., 2009). Indeed, the persistence of hexanucleotide motif-specific DNA methylation loss in cells completely devoid of TET activity, as well as in cells in which DNMT1 was the sole DNA methyltransferase present, strongly supports a DNMT1-centered mechanism. We note, however, that TET-deficient cells show considerably greater reduction of heterochromatic DNA methylation than control cells (López-Moyado et al., 2019). One explanation is simply that TET-deficient cells proliferate more than control cells in response to the same extrinsic signals (Moran-Crusio et al., 2011; Tsagaratou et al., 2017; Yang et al., 2025); the other is that TET deficiency feeds back to inhibit the activity of the DNMT1/UHRF1 complex in a manner that is still undefined. Further research will be necessary to distinguish these possibilities, which are not exclusive.

We observed that the hexanucleotide motif-specific DNMT1 preference and the proliferation-associated methylation loss are most pronounced in heterochromatic regions. These domains, which are often late-replicating and structurally constrained (Bell et al., 2023; Janssen et al., 2018), appear particularly sensitive to cumulative methylation maintenance errors. Importantly, methylation loss within heterochromatin is not uniform: individual CpGs show heterogeneous DNA methylation, even within regions that display overall hypomethylation. This heterogeneity argues against purely regional or chromatin-state–driven mechanisms, and instead highlights local, sequence-encoded determinants of DNA methylation stability.

These observations support a model in which methylation loss in phenomena such as aging and cancer—especially in heterochromatin—is not primarily driven by a regulated demethylation program but instead emerges from replication-associated imperfections in DNMT1-mediated maintenance. In physiological contexts, however, the DNMT1 intrinsic sequence biases are likely modulated by additional factors, including the activity of *de novo* methyltransferases such as DNMT3A and DNMT3B, as well as TET enzymes. Moreover, it is plausible that in certain biological phenomena, such as in cancer, selective pressures may further amplify these intrinsic DNMT1 biases, favoring retention or loss of methylation at specific motifs depending on their phenotypic consequences, thereby contributing to the pronounced methylation heterogeneity observed in cancer genomes (Brocks et al., 2014).

Noteworthy, the effects of the hexanucleotide DNA sequence context are probabilistic rather than deterministic. CpGs in low-ranking hexanucleotide motifs are not inevitably demethylated; rather, they have a higher probability of “not being remethylated” during any given cell cycle. Over many rounds of replication, these small biases accumulate, producing the reproducible, hexanucleotide motif-specific patterns of methylation loss that we observed in this study. This stochastic-yet-biased process provides a mechanistic explanation for both the emergence of DNA methylation heterogeneity with age and the selective vulnerability of specific CpGs to DNA hypomethylation during extensive cell proliferation.

The motif-specific hierarchy described here explains why DNA methylation loss during aging and proliferation is neither uniform nor random, but instead follows reproducible, sequence-dependent patterns. The reproducibility in DNA methylation loss is also a hallmark of DNA methylation-based aging clocks (Lu et al., 2023). According to our analysis, cell proliferation leaves an epigenetic record: each round of DNA replication introduces a probabilistic opportunity for methylation loss at low-ranking “vulnerable” CpGs, and over time these events accumulate into measurable, genome-wide hypomethylation patterns. Because our proposed hexanucleotide motif ranking reflects intrinsic DNMT1 maintenance efficiency, motif-specific methylation loss provides a quantitative readout of cumulative cell divisions and biological aging (**Fig. 5**). Our work also highlights that future DNA methylation clocks could be improved by explicitly incorporating the surrounding DNA sequence context, rather than treating CpGs as isolated indicators of DNA methylation changes.

Additionally, we propose that genomic regions requiring stable DNA methylation—such as imprinting control regions, or elements critical for genome integrity, such as transposable elements—have been subject to evolutionary or functional selection for high-ranking hexanucleotide motifs, in which DNA methylation will be maintained with higher fidelity. Consistent with this idea, high-ranking motifs are enriched in regulatory regions where methylation must be robustly preserved, suggesting that sequence composition itself contributes to protecting essential methylation states from cumulative maintenance errors. For instance, Krüppel-associated box-containing zinc finger protein (KRAB-ZFP) ZFP57, implicated in the establishment and maintenance of imprinted loci, binds to a hexanucleotide motif in a methylation-dependent manner (Quenneville et al., 2011), and its consensus binding site (T<u>GCCGCA</u>) includes a sequence (GCCGCA, ranked #2) that according to our ranking, is preferentially remethylated after replication. Similarly, the transcription factor YY1, implicated in paternal expression of the imprinted gene *Peg3* (Kim et al., 2003) and in transcription of LINE-1 elements in mouse and human cells (Athanikar et al., 2004; Saha et al., 2024), binds a motif containing the hexanucleotide GCCGCC (motif #47 of 225 in our ranking) (Kim & Kim, 2009). YY1 binding is inhibited by CpG methylation—YY1-binding sites in *Peg3* are methylated only on the maternal allele (Kim et al., 2003). Accordingly, methylation of this motif is likely precisely maintained, reflecting mechanisms that safeguard DNA methylation at critical motifs from maintenance errors and stochastic loss.

Conversely, the onset of proliferation in both normal and cancer cells is almost invariably accompanied by MYC upregulation or amplification (Dhanasekaran et al., 2022), and the canonical MYC E-box motif (CACGTG) (Blackwood et al., 1992; Prendergast et al., 1991) falls among the DNMT1 sequence motifs that are more susceptible to methylation loss during proliferation (motif #201 out of 225). Again, methylation of the CpG within the E-box motif inhibits MYC binding (Prendergast et al., 1991). Thus, transcription factor binding sites may be shaped not only by the underlying DNA sequence but also by motif-dependent methylation dynamics, emphasizing the still poorly appreciated point that transcription factor occupancy is shaped not only by motif identity but also by methylation state (Yin et al., 2017).

Collectively, our results demonstrate that differential DNA methylation loss across hexanucleotide CpG motifs is largely replication-dependent and can be attributed to imperfect maintenance of CpG methylation. The efficiency of this process is modulated by the flanking dinucleotide context, potentially reflecting a structural preference of the DNMT1-UHRF1 maintenance machinery for certain sequence contexts. This mechanism appears to be universal across cell types as well as the biological conditions associated with replication. Our analyses highlight a fundamental role for the flanking DNA sequences surrounding CpGs in shaping DNA methylation landscapes genome-wide.

## Methods

### Acute simultaneous deletion of Tet1, Tet2, and Tet3 genes in hematopoietic cells

*Tet1^fl/fl^/Tet2^fl/fl^/Tet3^fl/fl^ Rosa26-stop-EYFP UBC-Cre-ERT2* mice (*Tet iTKO* mice) have been described previously (Yuita et al., 2023). To delete floxed alleles using the *Cre-ERT2* recombinase, tamoxifen (Sigma) was solubilized at 10 mg mL^−1^ in corn oil (Sigma) and delivered into mice by intraperitoneal injection of 2 mg tamoxifen per mouse every day for five consecutive days. The day of the last tamoxifen injection was designated Day 0. Bone marrow cells were isolated after ∼28 days of tamoxifen treatment from *Tet iTKO* mice or control mice (*Rosa26-stop-EYFP UBC-Cre-ERT2* mice). All animal procedures were approved by the La Jolla Institute (LJI) Institutional Animal Care and Use Committee and were conducted in accordance with institutional guidelines.

### Myeloid cell sorting

Antibody staining and flow cytometric analysis was performed as previously described (Yuita et al., 2023). Briefly, bone marrow cells were flushed out of femurs and tibiae in Phosphate-Buffered Saline buffer (PBS, ThermoFisher Scientific) supplemented with 2% FBS + 10 mM HEPES. Cells were stained in FACS buffer (1% bovine serum albumin, 1 mM EDTA, and 0.05% sodium azide in PBS) with indicated antibodies for 30 mins on ice. Antibodies were purchased from BioLegend. The following monoclonal antibodies conjugated with allophycocyanin (APC) and phycoerythrin (PE) were used: Mac-1/CD11b (M1/70), Gr-1 (RB6-8C5). Efficient gene deletion was analyzed by YFP positivity. Cell sorting was performed using the BD FACSAria II, BD FACSAria Fusion, and the BD Influx Flow Cytometry Sorters (BD Biosciences), sorting for Cd11b^+^ Gr1^+^ YFP^+^ cells (hereafter referred to as myeloid cells).

### ONT-seq

After cell sorting, high-molecular-weight genomic DNA was isolated from *Tet iTKO* and *Control* myeloid cells using Nanobind CBB Kit (Circulomics, NB-900-001-01) according to the manufacturer’s instructions. DNA libraries were prepared at the Kinghorn Centre for Clinical Genomics (Australia) using 3 μg input DNA, without shearing, and an SQK-LSK109 ligation sequencing kit. Libraries were each sequenced separately on a PromethION (Oxford Nanopore Technologies) flow cell (FLO-PRO002, R9.4.1 chemistry). Bases were called with guppy 5.0.13 (Oxford Nanopore Technologies). Reads were aligned to the reference assembly using minimap2 2.17 and parameters -ax map-ont -L -t 64. CpG methylation calls were generated from ONT reads using nanopolish version 0.13.2118. The median coverage for each sample was ∼20x.

### Simultaneous detection of 5hmC and 5mC by duet multiomics solution evoC (biomodal)

After cell sorting, DNA was isolated from *Tet iTKO* and *Control* myeloid cells using the DNeasy Blood and Tissue kit (QIAGEN), according to the manufacturer’s instructions. The duet multiomics solution evoC method (6-base sequencing) (Füllgrabe et al., 2023) was performed according to the manufacturer’s instructions, using 70 ng of genomic DNA as starting material from *Tet iTKO* and *Control* myeloid cells. For 6-base sequencing, FASTQ files were processed using biomodal’s duet pipeline (version 1.4.1), as described in (Füllgrabe et al., 2023), and the median coverage for each sample was ∼19x.

### DNA methylation analyses

For the reanalyzed datasets, we employed Bismark (v0.25.1) (Krueger & Andrews, 2011) to align reads from bisulfite-treated samples to either the mm10 mouse reference genome or the hg38 human reference genome using default parameters. Single and paired-end reads were mapped as appropriately. Bismark methylation extractor (v0.25.1) (Krueger & Andrews, 2011) was employed to estimate CpG methylation from the aligned reads and methylation ratios on both DNA strands were merged for each CpG site. Only CpG sites covered by at least 5 reads (5x coverage) and within the autosomes were considered for the analysis (chrX and chrY were excluded from the analyses). For euchromatin/heterochromatin genome compartmentalization, principal component analysis (PCA) of Hi-C data (A/B compartment identification) was performed as described in our previous publication (López-Moyado et al., 2019). List of previously published DNA methylation datasets is provided in **Supplementary Table 1**.

### Hexanucleotide motif analysis

Hexanucleotide motifs were obtained for each covered CpG by extracting two nucleotides upstream and three nucleotides downstream of the cytosine. The genomic coordinates of the covered cytosines were derived from the Bismark methylation extractor (v0.25.1) (Krueger & Andrews, 2011) output. Only CpGs with sufficient coverage (≥5x) were considered in the analysis. The coordinates of these CpGs were expanded to cover the hexanucleotide context in R (v4.4.1) using the Bioconductor packages BSgenome (v1.72.1), Biostrings (v2.72.1), GenomicRanges(v1.56.2), rtracklayer (v1.64.0) and plyranges (v1.24.0). The corresponding sequences were extracted using the same tools together with the appropriate reference genome for each dataset (e.g., BSgenome.Mmusculus.UCSC.mm10 (v1.4.3), BSgenome.Hsapiens.UCSC.hg38 (v1.4.5)). For genome browser visualizations, tracks for specific hexanucleotide contexts were generated in R (v4.4.1) and visualized using the Integrative Genomics Viewer (IGV) (v3.0.1).

### Statistical analyses

Statistical analyses and bar plots were performed and plotted with R (v4.4.1). The bar graph and dot plots shown indicate mean and SE. Most comparisons were analyzed using two-tailed unpaired *t* test, as indicated in the figure legends unless otherwise stated. LOESS fitted regression lines plus Spearman correlation scores were calculated for each DNA methylation difference scatter plot, as indicated in the figure legends unless otherwise stated. Asterisks indicate statistically significant differences: \*\*\*\**P*< 0.0001, \*\*\**P*< 0.001, \*\**P*< 0.01, \**P*<0.05.

## Data availability

GEO accession codes for the previously published sequencing datasets used in this study are available in **Supplementary Table 1**. Sequencing data produced as part of this study (ONT-seq, 6-letter-seq) have been deposited in the Gene Expression Omnibus (GEO) database under accession codes **GSE325385** (ONT-seq) and **GSE324863** (6-letter-seq).

## Acknowledgements

This publication includes data generated at the UCSD Human Embryonic Stem Cell Core Facility (UCSD HESCCF), using the BD FACS Aria Fusion, BD FACS Aria II, and the BD Influx Flow Cytometry Sorters, supported by the CIRM Major Facilities grant (FA1-00607) to the Sanford Consortium for Regenerative Medicine. We thank Cody Fine, Mateo Espinoza and Mitra Banihassan (UCSD HESCCF) for technical assistance with cell sorting experiments; and the Department of Laboratory Animal Care at La Jolla Institute for Immunology and the animal facility for excellent support. ONT sequencing was done using a PromethION at the Garvan Sequencing Platform in Sydney. This research was supported by NIH grants R35 CA210043 and U01 AI180152 to A.R., EvansMDS Young Investigator Award to I.F.L.-M., and NHMRC Investigator Grant (GNT2033366) and the Mater Foundation to G.J.F.

## Author contributions

I.F.L.-M., L.H.-E., and A.R. designed research and analyzed data; I.F.L.-M., L.H.-E., and J.C.A. performed bioinformatic analyses; I.F.L.-M. performed mouse experiments, myeloid cell isolation, and DNA extraction; A.M., E.L., and R.C. prepared the libraries and processed the 6-letter-seq datasets; G.J.F. performed and processed ONT sequencing; and I.F.L.-M. and A.R. wrote the manuscript with input from all the authors.

## Supplementary materials

Supplementary figures S1-S6

Supplementary tables S1-S2

Supplementary table S1. List of previously published DNA methylation datasets used in this study.

Supplementary table S2. Numbered rank list of hexanucleotide CpG motifs.

