## Supplementary Materials for "Flanking DNA sequences determine DNA methylation maintenance in proliferation, cancer and aging"

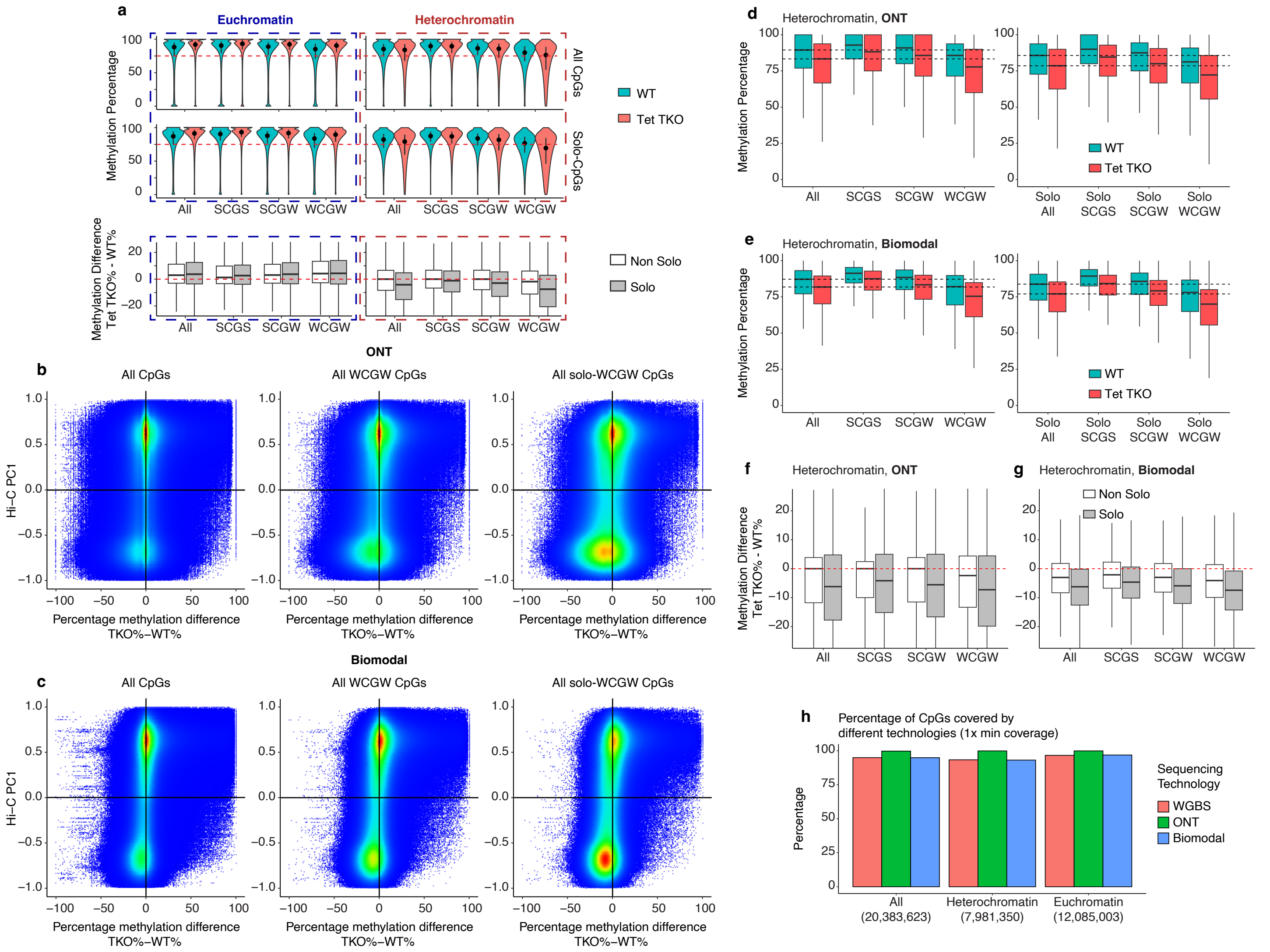


**Supplemental Figure S1: Solo-WCGW CpGs display the greatest susceptibility to loss of DNA methylation in *Tet* triple deficient cells.**

**(a)** DNA methylation percentage (*top*) and methylation difference (*bottom*) in *Tet1/2/3 TKO* myeloid cells across all or different sequence contexts (SCGS, SCGW, WCGW) in euchromatin and heterochromatin compartments, as measured by WGBS.

**(b)** Percentage DNA methylation difference (*x-axis*) in euchromatin/heterochromatin as determined by Hi-C PC1 values (*y-axis*) in *Tet1/2/3 TKO* myeloid cells (isolated ex vivo after ~28 days of tamoxifen treatment of *Tet1/2/3 triple-floxed CreERT2* mice), across all CpGs, those in the WCGW context, and those in CpG-poor regions (*solo*, no other CpGs +/- 35 bp), as measured by ONT sequencing.

**(c)** Percentage DNA methylation difference (*x-axis*) in euchromatin/heterochromatin as determined by Hi-C PC1 values (*y-axis*, positive values represent euchromatin, negative values represent heterochromatin) in *Tet1/2/3 TKO* myeloid cells (isolated ex vivo after ~28 days of tamoxifen treatment of *Tet1/2/3 triple-floxed CreERT2* mice), across all CpGs, those in the WCGW context, and those in CpG-poor regions (*solo*, no other CpGs +/- 35 bp), as measured by 6-base-seq sequencing.

**(d, f)** DNA methylation percentage (**d**) and methylation difference (**f**) in *Tet1/2/3 TKO* myeloid cells, across all or different sequence contexts (SCGS, SCGW, WCGW) and CpG densities (*solo* defined as no other CpGs +/- 35 bp) in heterochromatin, as measured by ONT sequencing.

**(e, g)** DNA methylation percentage (**e**) and methylation difference (**g**) in *Tet1/2/3 TKO* myeloid cells, across all or different sequence contexts (SCGS, SCGW, WCGW) and CpG densities (*solo* defined as no other CpGs +/- 35 bp) in heterochromatin, as measured by 6-base-seq sequencing.

**(h)** Percentage of CpGs covered by WGBS, ONT, and 6-base-seq technologies across the genome, and euchromatin and heterochromatin compartments.

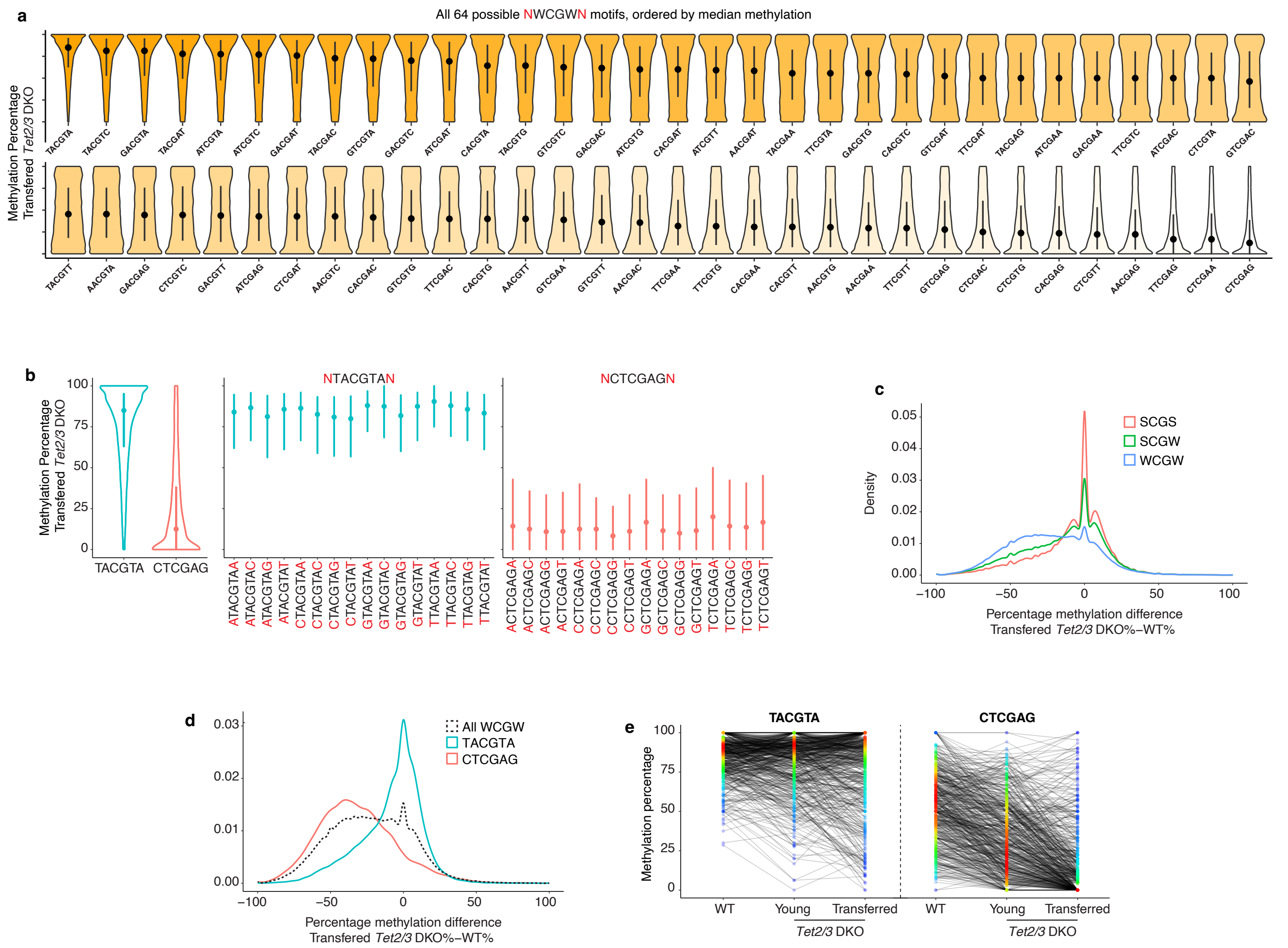


**Supplemental Figure S2: Solo-WCGW CpGs in different hexanucleotide sequence contexts display distinct DNA methylation levels and a graded susceptibility to loss of DNA methylation.**

**(a)** DNA methylation percentage in *Tet2/3 DKO* transferred iNKT cells, across all 64 possible hexanucleotide CpG sequences found in the solo-WCGW context in heterochromatin, ordered and colored from highest to lowest median DNA methylation per hexanucleotide sequence. Dots represent the median and lines the quartiles for each sequence. TACGTA and CTCGAG sequences appear in the first and last position, respectively.

**(b)** DNA methylation percentage in *Tet2/3 DKO* transferred iNKT cells, for the heterochromatic CpGs in the TACGTA and CTCGAG context (left), and across all the possible 8-nucleotide CpG sequences found in the NTACGTAN (middle) and NCTCGAGN (right) sequence contexts. Dots represent the median and lines the quartiles for each sequence. NTACGTAN and NCTCGAGN sequences are highlighted in blue and orange, respectively.

**(c)** Density distribution of the difference in DNA methylation percentage between *Tet2/3 DKO* transferred iNKT cells and control iNKT cells in different flanking contexts for heterochromatic solo-CpGs.

**(d)** Density distribution of the difference in DNA methylation percentage between *Tet2/3 DKO* transferred iNKT cells and control iNKT cells in TACGTA and CTCGAG flanking contexts for heterochromatic solo-CpGs.

**(e)** DNA methylation percentage of heterochromatic (Hi-C PC1 value < 0) solo-WCGW CpGs in WT, *Tet2/3 DKO* young, and *Tet2/3 DKO* transferred iNKT cells, found within the TACGTA context (left) and within the CTCGAG context (right).

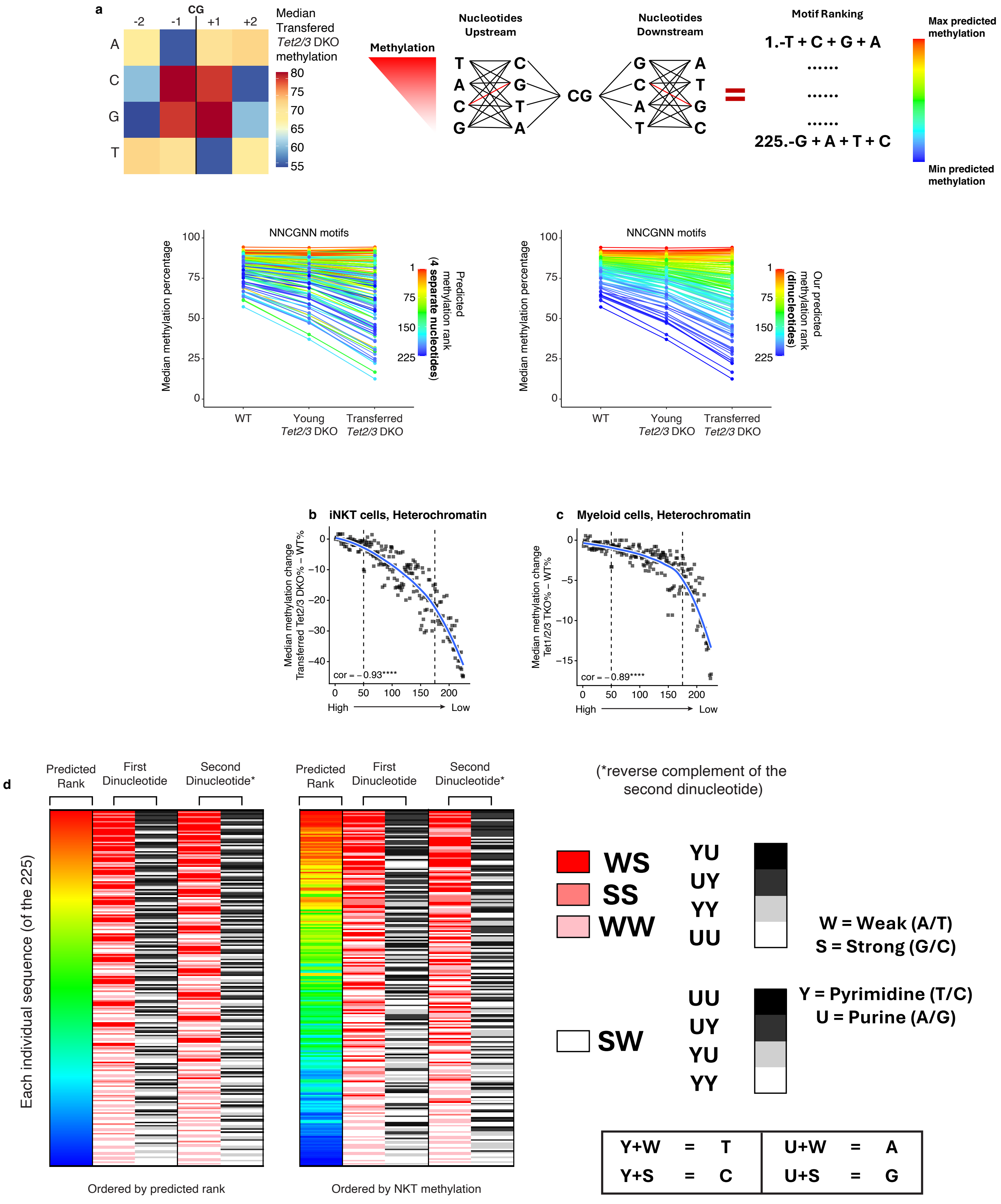


**Supplemental Figure S3: The hexanucleotide sequence context surrounding a CpG determines its DNA methylation level and susceptibility to lose DNA methylation.**

**(a)** *Top*, Ranking CpG methylation in WT, *Tet2/3 DKO* young, and *Tet2/3 DKO* transferred iNKT cells by the effects of individual 5' and 3' nucleotides at the -1 and +1 positions (*top left*), compared to dinucleotides (combination of -1/-2 and +1/+2 positions, *top right* and **Fig. 3i**). *Bottom*, Median DNA methylation percentage in WT, *Tet2/3 DKO* young, and *Tet2/3 DKO* transferred iNKT cells, across all 225 possible 6-nucleotide CpG sequences found in the *solo* context in heterochromatin, colored by predicted methylation rank based on individual flanking nucleotides (*left*) and our proposed dinucleotide hierarchy (*right*). AR: Let me know if I have this right; can you also put a complete numbered rank list in a suppl table?

**(b)** Correlation between hexanucleotide CpG motif ranking and median DNA methylation difference between WT and *Tet2/3 DKO* transferred iNKT cells, in heterochromatic solo-CpGs. Fitted regression lines plus Spearman correlation scores for each DNA methylation difference scatter plot are shown (*bottom left corner*). **** denotes $p < 0.0001$ (Spearman correlation).

**(c)** Correlation between hexanucleotide CpG motif ranking and median DNA methylation difference between WT and *Tet1/2/3 TKO* ex vivo myeloid cells, in heterochromatic solo-CpGs. Fitted regression lines plus Spearman correlation scores for each DNA methylation difference scatter plot are shown (*bottom left corner*). **** denotes $p < 0.0001$ (Spearman correlation).

**(d)** DNA methylation rank is determined by the combination of features in 5’ and 3’ dinucleotides: whether pyrimidines or purines, and whether they engage in strong (C:G, termed S) or weak (A:T, termed W) base pairing.

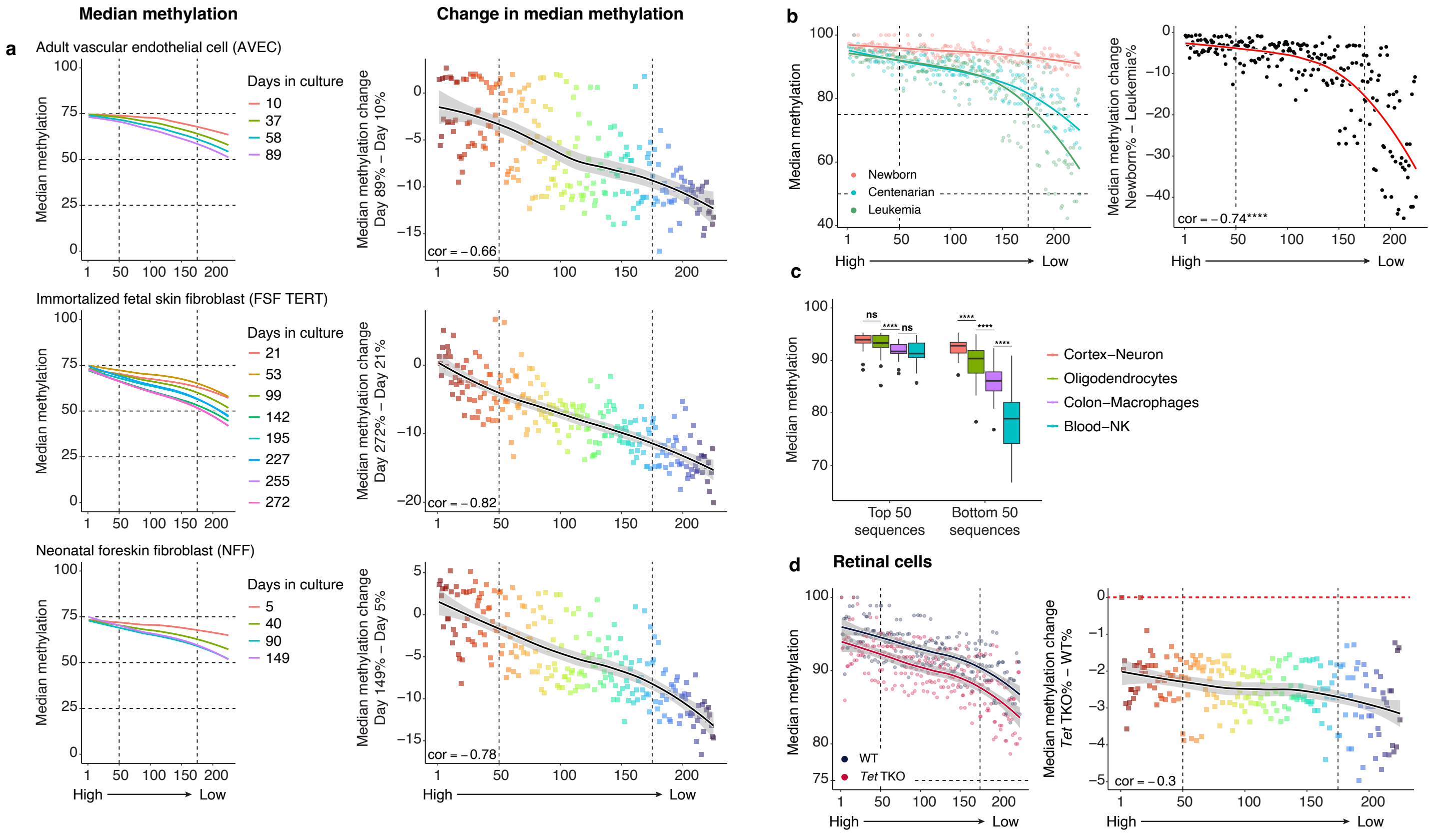


**Supplemental Figure S4: Loss of DNA methylation in lowly ranked motifs in influenced by cell proliferation.**

(**a**) Genome-wide correlation between hexanucleotide CpG motif ranking and median DNA methylation percentage for adult vascular endothelial cells (AVEC, *top*), TERT-immortalized foreskin skin fibroblasts (FSF TERT, *middle*) and NFF (neonatal foreskin fibroblasts, *bottom*). *Left*, data for different days in culture; *right*, median DNA methylation difference between the last and first day.

(**b**) *Left*, Genome-wide correlation between hexanucleotide CpG motif ranking and median DNA methylation percentage for $CD4^+$ T cells from a newborn, a centenarian and a T cell leukemia. *Right*, median DNA methylation difference between newborn and leukemic T cells.

(**c**) Median DNA methylation percentage for the top 50 sequences (ranked 1-50) and bottom 50 sequences (ranked 175-225) in two primarily post-mitotic cell types (cortex neurons and oligodendrocytes), and two rapidly proliferating cell types (Natural Killer cells and colon macrophages). **** denotes $p < 0.0001$ (*t*-test).

(**d**) Genome-wide correlation between hexanucleotide CpG motif ranking and either median DNA methylation percentage (*left*) and median DNA methylation difference (*right*) in *TET TKO* retinal cells, which are largely post-mitotic. Fitted regression lines along their confidence intervals are displayed for each scatter plot.

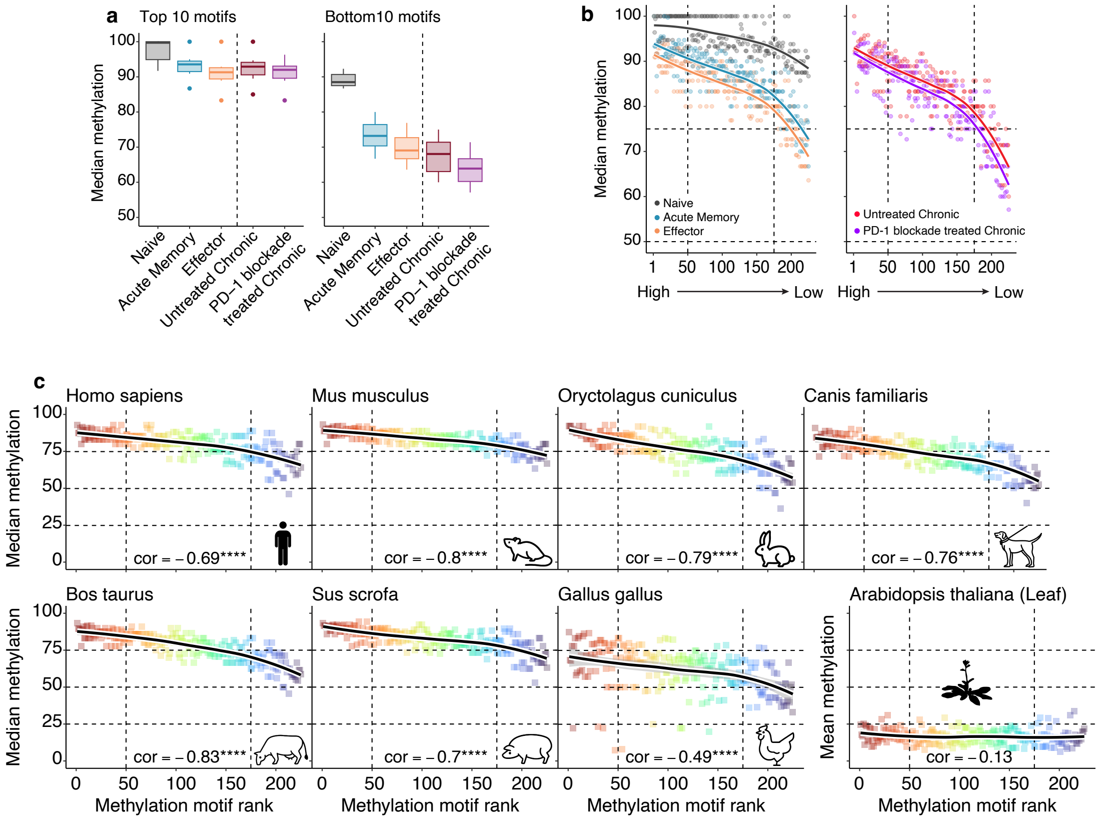


**Supplemental Figure S5: Loss of DNA methylation at low-ranking CpG motifs is associated with $CD8^+$ T cell subsets that have undergone greater proliferation and the hexanucleotide ranking is evolutionarily conserved in vertebrates.**

(**a**) Median DNA methylation percentage for the top 10 sequences/motifs (ranked 1-10) and bottom 50 sequences (ranked 216-225) in different subsets from $CD8^+$ T cells from mice exposed to chronic or acute LCMV infection, or from chronically infected mice treated with anti-PD-1.

(**b**) Genome-wide correlation between hexanucleotide CpG motif ranking and median DNA methylation percentage in different $CD8^+$ T cell subsets from mice exposed to chronic or acute LCMV infection, or from chronically infected mice treated with anti-PD-1.

(**c**) Genome-wide correlation between hexanucleotide CpG motif ranking and median DNA methylation percentage in fibroblasts from 7 different vertebrate species; data from leaves of *Arabidopsis thaliana* are shown for comparison. Fitted regression lines along their confidence intervals plus Spearman correlation scores for each DNA methylation scatter plot are shown. **** denotes $p < 0.0001$ (Spearman correlation).

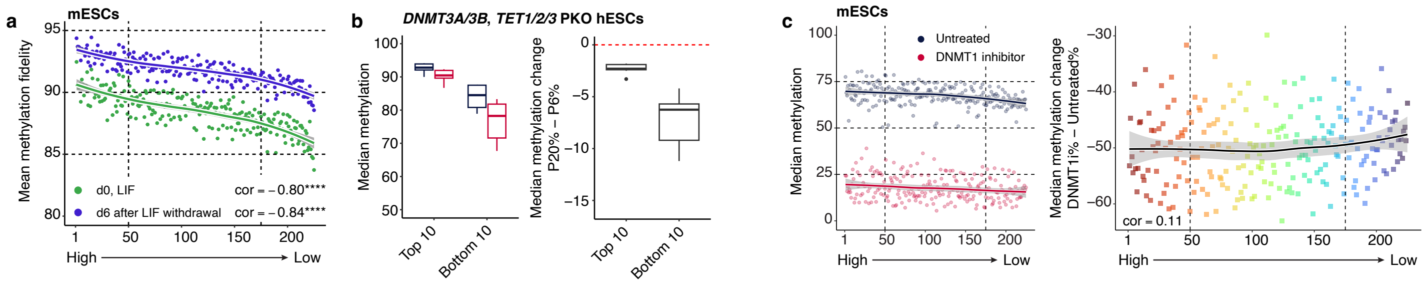


**Supplemental Figure S6: Loss of DNA methylation associated with imperfect DNMT1 activity at disfavored CpG hexanucleotide motifs.**

(**a**) Genome-wide correlation between hexanucleotide CpG motif ranking and mean DNA methylation fidelity (defined as the percentage of symmetrically methylated CpGs with respect to total CpGs—total CpGs defined as the sum of symmetrically methylated, symmetrically unmethylated and hemimethylated CpGs), as assayed in undifferentiated mESC at day 0 and upon differentiation 6 days after LIF withdrawal. Fitted regression lines along their confidence intervals plus Spearman correlation scores for each mean DNA methylation fidelity scatter plot are shown. **** denotes $p < 0.0001$ (Spearman correlation).

(**b**) Median DNA methylation percentage (*left*) and median DNA methylation difference (*right*) for the top 10 sequences/motifs (#1-10) and bottom 10 sequences (# 216-225) in hESC expressing only DNMT1 (from **Fig. 6a**).

(**c**) Genome-wide correlation between 6-nucleotide CpG motif ranking and median DNA methylation percentage (*left*), or median DNA methylation difference (*right*) in mESCs after 14 days of GSK-3484862 treatment. Fitted regression lines along their confidence intervals are displayed for each scatter plot.

**Supplementary table S1. List of previously published DNA methylation datasets used in this study.**

| Sample | Sample type | Accession Number | PMID | Organism | Genome Assembly |
|---|---|---|---|---|---|
| NKT | WT<br>Young Tet 2/3 DKO | GSE66834 | 27869820 | Mus musculus | mm10 |
| | Transferred Tet 2/3 DKO | GSE134396 | 31371502 | | |
| Myeloid CD11b+ | WT<br>Tet 1/2/3 TKO | GSE222726 | 37406303 | Mus musculus | mm10 |
| Primary human cell cultures | Adult vascular endothelial cells<br>Immortalized fetal skin fibroblasts<br>Neonatal foreskin fibroblasts | GSE197545 | 36347867 | Homo sapiens | hg38 |
| Retinal neurons | WT<br>Tet 1/2/3 TKO | PRJNA1121391 | 40053367 | Mus musculus | mm10 |
| CD4+T Cells | Newborn<br>103 year old | GSE31263 | 22689993 | Homo sapiens | hg38 (realigned) |
| Colorectal tissue | Normal colonic mucosa<br>CIMP-H colon tumor | GSE32399 | 22120008 | Homo sapiens | hg18 |
| IMR-90 | Proliferating<br>Senescent | GSE48580 | 27457071 | Homo sapiens | hg18 |
| Diverse human cell types | … | GSE186458 | 36599988 | Homo sapiens | hg38 |
| CD8+T Cells | Aged naïve<br>Young memory<br>0.5 Lifetimes<br>2 Lifetimes d1350<br>2 Lifetimes d1620<br>4 Lifetimes | GSE263941 | 38867059 | Mus musculus | mm10 |
| CD4+T Cells | 25yo indiv3<br>82yo indiv4<br>82yo indiv5<br>86yo indiv6 | GSE86340 | 28346445 | Homo sapiens | hg38 (realigned) |
| CD4+T Cells | Prolymphocytic Leukemia | S016KWU1 (BLUEPRINT) | 29610480 | Homo sapiens | hg38 (realigned) |
| CD8+T Cells | Naïve<br>Acute Memory<br>Effector<br>Untreated Chronic<br>PD-1 blockade treated Chronic | GSE99451 | 28648661 | Mus musculus | mm10 |
| E14 mESCs | d0, with LIF<br>d6 after LIF withdrawal | GSE48229 | 24835587 | Mus musculus | mm10 |
| hESCs Dnmt3a/b Tet1/2/3 PKO | Passage 6<br>Passage 20 | GSE126958 | 32514123 | Homo sapiens | hg19 |
| mESCs Dnmt3a/b DKO + shDnmt1 | Passage 1 after Dnmt1 reconstitution<br>Passage 5 after Dnmt1 reconstitution<br>Passage 15 after Dnmt1 reconstitution<br>Passage 25 after Dnmt1 reconstitution | GSE158460 | 34140676 | Mus musculus | mm9 |
| mESCs Dnmt1/3a/3b TKO | No reconstitution<br>full-length Dnmt1 reconstitution<br>602 mutant Dnmt1 reconstitution | GSE200166 | 38811355 | Mus musculus | mm10 |
| mESCs | DMSO day 14<br>2uM GSK-3484862 d14 | GSE184116 | 34906184 | Mus musculus | mm10 |
| mESCs | WT<br>Ogt KO | GSE247534 | 40155743 | Mus musculus | mm10 |
| Fibroblasts | | GSE175615 | 35317801 | Homo sapiens<br>Mus musculus<br>Oryctolagus cuniculus<br>Canis lupus familiaris<br>Bos taurus<br>Sus scrofa<br>Gallus gallus | hg38<br>mm10<br>oryCun2<br>canFam3<br>bosTau8<br>susScr11<br>galGal6 |
| Leaf | WT A Rosette leaf | GSE191006 | 34954804 | Arabidopsis thaliana | Araport11 |

**Supplementary table S2. Numbered rank list of hexanucleotide CpG motifs.**

| Rank | Hexanucleotide_motif | 5' dinucleotide | 3' dinucleotide |
|---|---|---|---|
| 1 | TGCGCA | TG | CA |
| 2 | TGCGGC | TG | GC |
| 3 | GCCGCA | GC | CA |
| 4 | TGCGGT | TG | GT |
| 5 | GCCGGC | GC | GC |
| 6 | ACCGCA | AC | CA |
| 7 | TGCGGA | TG | GA |
| 8 | GCCGGT | GC | GT |
| 9 | ACCGGC | AC | GC |
| 10 | TCCGCA | TC | CA |
| 11 | TGCGCT | TG | CT |
| 12 | GCCGGA | GC | GA |
| 13 | ACCGGT | AC | GT |
| 14 | TCCGGC | TC | GC |
| 15 | AGCGCA | AG | CA |
| 16 | TGCGTA | TG | TA |
| 17 | GCCGCT | GC | CT |
| 18 | ACCGGA | AC | GA |
| 19 | TCCGGT | TC | GT |
| 20 | AGCGGC | AG | GC |
| 21 | TACGCA | TA | CA |
| 22 | TGCGGG | TG | GG |
| 23 | GCCGTA | GC | TA |
| 24 | ACCGCT | AC | CT |
| 25 | TCCGGA | TC | GA |
| 26 | AGCGGT | AG | GT |
| 27 | TACGGC | TA | GC |
| 28 | CCCGCA | CC | CA |
| 29 | TGCGAT | TG | AT |
| 30 | GCCGGG | GC | GG |
| 31 | ACCGTA | AC | TA |
| 32 | TCCGCT | TC | CT |
| 33 | AGCGGA | AG | GA |
| 34 | TACGGT | TA | GT |
| 35 | CCCGGC | CC | GC |
| 36 | ATCGCA | AT | CA |
| 37 | TGCGCC | TG | CC |
| 38 | GCCGAT | GC | AT |
| 39 | ACCGGG | AC | GG |
| 40 | TCCGTA | TC | TA |
| 41 | AGCGCT | AG | CT |
| 42 | TACGGA | TA | GA |
| 43 | CCCGGT | CC | GT |
| 44 | ATCGGC | AT | GC |
| 45 | GGCGCA | GG | CA |
| 46 | TGCGTC | TG | TC |
| 47 | GCCGCC | GC | CC |
| 48 | ACCGAT | AC | AT |
| 49 | TCCGGG | TC | GG |
| 50 | AGCGTA | AG | TA |
| 51 | TACGCT | TA | CT |
| 52 | CCCGGA | CC | GA |
| 53 | ATCGGT | AT | GT |
| 54 | GGCGGC | GG | GC |

| | | | |
|---|---|---|---|
| 55 | GACGCA | GA | CA |
| 56 | TGCGAC | TG | AC |
| 57 | GCCGTC | GC | TC |
| 58 | ACCGCC | AC | CC |
| 59 | TCCGAT | TC | AT |
| 60 | AGCGGG | AG | GG |
| 61 | TACGTA | TA | TA |
| 62 | CCCGCT | CC | CT |
| 63 | ATCGGA | AT | GA |
| 64 | GGCGGT | GG | GT |
| 65 | GACGGC | GA | GC |
| 66 | GTCGCA | GT | CA |
| 67 | TGCGTG | TG | TG |
| 68 | GCCGAC | GC | AC |
| 69 | ACCGTC | AC | TC |
| 70 | TCCGCC | TC | CC |
| 71 | AGCGAT | AG | AT |
| 72 | TACGGG | TA | GG |
| 73 | CCCGTA | CC | TA |
| 74 | ATCGCT | AT | CT |
| 75 | GGCGGA | GG | GA |
| 76 | GACGGT | GA | GT |
| 77 | GTCGGC | GT | GC |
| 78 | CACGCA | CA | CA |
| 79 | TGCGTT | TG | TT |
| 80 | GCCGTG | GC | TG |
| 81 | ACCGAC | AC | AC |
| 82 | TCCGTC | TC | TC |
| 83 | AGCGCC | AG | CC |
| 84 | TACGAT | TA | AT |
| 85 | CCCGGG | CC | GG |
| 86 | ATCGTA | AT | TA |
| 87 | GGCGCT | GG | CT |
| 88 | GACGGA | GA | GA |
| 89 | GTCGGT | GT | GT |
| 90 | CACGGC | CA | GC |
| 91 | AACGCA | AA | CA |
| 92 | TGCGAA | TG | AA |
| 93 | GCCGTT | GC | TT |
| 94 | ACCGTG | AC | TG |
| 95 | TCCGAC | TC | AC |
| 96 | AGCGTC | AG | TC |
| 97 | TACGCC | TA | CC |
| 98 | CCCGAT | CC | AT |
| 99 | ATCGGG | AT | GG |
| 100 | GGCGTA | GG | TA |
| 101 | GACGCT | GA | CT |
| 102 | GTCGGA | GT | GA |
| 103 | CACGGT | CA | GT |
| 104 | AACGGC | AA | GC |
| 105 | TTCGCA | TT | CA |
| 106 | TGCGAG | TG | AG |
| 107 | GCCGAA | GC | AA |
| 108 | ACCGTT | AC | TT |
| 109 | TCCGTG | TC | TG |
| 110 | AGCGAC | AG | AC |
| 111 | TACGTC | TA | TC |

| | | | |
|---|---|---|---|
| 112 | CCCGCC | CC | CC |
| 113 | ATCGAT | AT | AT |
| 114 | GGCGGG | GG | GG |
| 115 | GACGTA | GA | TA |
| 116 | GTCGCT | GT | CT |
| 117 | CACGGA | CA | GA |
| 118 | AACGGT | AA | GT |
| 119 | TTCGGC | TT | GC |
| 120 | CTCGCA | CT | CA |
| 121 | GCCGAG | GC | AG |
| 122 | ACCGAA | AC | AA |
| 123 | TCCGTT | TC | TT |
| 124 | AGCGTG | AG | TG |
| 125 | TACGAC | TA | AC |
| 126 | CCCGTC | CC | TC |
| 127 | ATCGCC | AT | CC |
| 128 | GGCGAT | GG | AT |
| 129 | GACGGG | GA | GG |
| 130 | GTCGTA | GT | TA |
| 131 | CACGCT | CA | CT |
| 132 | AACGGA | AA | GA |
| 133 | TTCGGT | TT | GT |
| 134 | CTCGGC | CT | GC |
| 135 | ACCGAG | AC | AG |
| 136 | TCCGAA | TC | AA |
| 137 | AGCGTT | AG | TT |
| 138 | TACGTG | TA | TG |
| 139 | CCCGAC | CC | AC |
| 140 | ATCGTC | AT | TC |
| 141 | GGCGCC | GG | CC |
| 142 | GACGAT | GA | AT |
| 143 | GTCGGG | GT | GG |
| 144 | CACGTA | CA | TA |
| 145 | AACGCT | AA | CT |
| 146 | TTCGGA | TT | GA |
| 147 | CTCGGT | CT | GT |
| 148 | TCCGAG | TC | AG |
| 149 | AGCGAA | AG | AA |
| 150 | TACGTT | TA | TT |
| 151 | CCCGTG | CC | TG |
| 152 | ATCGAC | AT | AC |
| 153 | GGCGTC | GG | TC |
| 154 | GACGCC | GA | CC |
| 155 | GTCGAT | GT | AT |
| 156 | CACGGG | CA | GG |
| 157 | AACGTA | AA | TA |
| 158 | TTCGCT | TT | CT |
| 159 | CTCGGA | CT | GA |
| 160 | AGCGAG | AG | AG |
| 161 | TACGAA | TA | AA |
| 162 | CCCGTT | CC | TT |
| 163 | ATCGTG | AT | TG |
| 164 | GGCGAC | GG | AC |
| 165 | GACGTC | GA | TC |
| 166 | GTCGCC | GT | CC |
| 167 | CACGAT | CA | AT |
| 168 | AACGGG | AA | GG |

| | | | |
|---|---|---|---|
| 169 | TTCGTA | TT | TA |
| 170 | CTCGCT | CT | CT |
| 171 | TACGAG | TA | AG |
| 172 | CCCGAA | CC | AA |
| 173 | ATCGTT | AT | TT |
| 174 | GGCGTG | GG | TG |
| 175 | GACGAC | GA | AC |
| 176 | GTCGTC | GT | TC |
| 177 | CACGCC | CA | CC |
| 178 | AACGAT | AA | AT |
| 179 | TTCGGG | TT | GG |
| 180 | CTCGTA | CT | TA |
| 181 | CCCGAG | CC | AG |
| 182 | ATCGAA | AT | AA |
| 183 | GGCGTT | GG | TT |
| 184 | GACGTG | GA | TG |
| 185 | GTCGAC | GT | AC |
| 186 | CACGTC | CA | TC |
| 187 | AACGCC | AA | CC |
| 188 | TTCGAT | TT | AT |
| 189 | CTCGGG | CT | GG |
| 190 | ATCGAG | AT | AG |
| 191 | GGCGAA | GG | AA |
| 192 | GACGTT | GA | TT |
| 193 | GTCGTG | GT | TG |
| 194 | CACGAC | CA | AC |
| 195 | AACGTC | AA | TC |
| 196 | TTCGCC | TT | CC |
| 197 | CTCGAT | CT | AT |
| 198 | GGCGAG | GG | AG |
| 199 | GACGAA | GA | AA |
| 200 | GTCGTT | GT | TT |
| 201 | CACGTG | CA | TG |
| 202 | AACGAC | AA | AC |
| 203 | TTCGTC | TT | TC |
| 204 | CTCGCC | CT | CC |
| 205 | GACGAG | GA | AG |
| 206 | GTCGAA | GT | AA |
| 207 | CACGTT | CA | TT |
| 208 | AACGTG | AA | TG |
| 209 | TTCGAC | TT | AC |
| 210 | CTCGTC | CT | TC |
| 211 | GTCGAG | GT | AG |
| 212 | CACGAA | CA | AA |
| 213 | AACGTT | AA | TT |
| 214 | TTCGTG | TT | TG |
| 215 | CTCGAC | CT | AC |
| 216 | CACGAG | CA | AG |
| 217 | AACGAA | AA | AA |
| 218 | TTCGTT | TT | TT |
| 219 | CTCGTG | CT | TG |
| 220 | AACGAG | AA | AG |
| 221 | TTCGAA | TT | AA |
| 222 | CTCGTT | CT | TT |
| 223 | TTCGAG | TT | AG |
| 224 | CTCGAA | CT | AA |
| 225 | CTCGAG | CT | AG |
